# PIP_2_-dependent coupling of voltage sensor and pore domains in K_v_7.2

**DOI:** 10.1101/2021.05.11.443644

**Authors:** Shashank Pant, Jiaren Zhang, Eung Chang Kim, Kin Lam, Hee Jung Chung, Emad Tajkhorshid

**Author notes:** **CORRESPONDING AUTHORS: Emad Tajkhorshid**, Beckman Institute for Advanced Science and Technology, University of Illinois at Urbana-Champaign, 405 N Mathews Avenue, 3147 Beckman Institute, Urbana, IL, 61801, USA., **Hee Jung Chung**, Department of Molecular and Integrative Physiology, University of Illinois at Urbana-Champaign, 407 South Goodwin Avenue, 524 Burrill Hall, Urbana, IL 61801, USA. These authors contributed equally (Shashank Pant, Jiaren Zhang, Eung Chang Kim). **Author Contributions** S.P., J.Z., H.J.C. and E.T. conceived of the study and participated in its design and coordination. S.P., K.L. and E.T. performed and analyzed MD simulation. E.C.K., J.Z. and H.J.C. generated and characterized selected mutations. S.P., J.Z., E.C.K., H.J.C., and E.T. wrote the manuscript. All authors read and approved the final manuscript.

## Abstract

Phosphatidylinositol-4,5-bisphosphate (PIP_2_) is a signaling lipid which regulates voltage-gated K_v_7/*KCNQ* potassium channels. Altered PIP_2_ sensitivity of neuronal K_v_7.2 channel is involved in *KCNQ2* epileptic encephalopathy. However, the molecular action of PIP_2_ on K_v_7.2 gating remains largely elusive. Here, we use molecular dynamics simulations and electrophysiology to characterize PIP_2_ binding sites in a human K_v_7.2 channel. In the closed state, PIP_2_ localizes to the periphery of the voltage-sensing domain (VSD). In the open state, PIP_2_ binds to 4 distinct interfaces formed by the cytoplasmic ends of the VSD, the gate, intracellular helices A and B and their linkers. PIP_2_ binding induces bilayer-interacting conformation of helices A and B and the correlated motion of the VSD and the pore domain, whereas charge-neutralizing mutations block this coupling and reduce PIP_2_ sensitivity of K_v_7.2 channels by disrupting PIP_2_ binding. These findings reveal the allosteric role of PIP_2_ in K_v_7.2 channel activation.

## Introduction

Phosphoinositides are major constituents of biological membranes and key regulators of fundamental biological processes including signal transduction, membrane trafficking, and cytoskeletal dynamics ^1^. Among the phosphoinositides, phosphatidylinositol-4,5-bisphosphate (PIP_2_) in the plasma membrane serves as a critical cofactor for many ion channels despite its low abundance (∼1% of total acidic lipids) ^2,3^. The affected channels include inward rectifier potassium (K^+^) channels ^4^, voltage-gated calcium channels, transient receptor potential channels, hyperpolarization-activated cyclic nucleotide-gated channels, and voltage-gated potassium (K_v_) channels ^2,5^.

PIP_2_ activates all five members of the K_v_ channel subfamily Q (K_v_7.1-K_v_7.5) which control excitability of neuronal, sensory, and muscle cells ^6,7^. Encoded by *KCNQ*1-*KCNQ*5 genes ^7^, each K_v_7 subunit has six transmembrane segments ^8,9^. The first four segments (S1-S4) comprise a voltage-sensing domain (VSD) with the S4 being the main voltage-sensor ^8,9^. The pore domain consists of the last two segments (S5-S6) flanking the pore loop which contains a highly conserved sequence and structure for K^+^ selectivity and permeability ^8,9^. The C-terminal intersection of four S6 segments constitutes the main gate ^9,10^. Each subunit also has a long intracellular C-terminal tail that harbors four helices (A-D) ^11^. Helix-A and Helix-B interact with calmodulin (CaM) ^11^, Helix-C mediates inter-subunit interaction, while Helix-D specifies the subunit assembly as a homotetramer or a heterotetramer ^11^.

Despite the common core structure, each K_v_7 subunit follows a distinct, cell-specific distribution that dictates its physiological roles in different tissues ^7,8^. In the heart, K_v_7.1 assembles with an auxiliary β subunit KCNE1 to produce the slow K^+^ current critical for repolarizing cardiac action potentials (APs) ^7,8^. K_v_7.4 is primarily found in cochlear hair cells of the inner ear ^8^. In the central nervous system, K_v_7 channels are mostly heterotetramers of K_v_7.2 and K_v_7.3, and to a lesser extent heterotetramers of K_v_7.3 and K_v_7.5 and homomeric K_v_7.2 channels ^12,13^. Neuronal K_v_7 channels produce slowly activating and non-inactivating K^+^ current (*I*_M_) that suppresses repetitive firing of APs ^12,13^, and dominant mutations in their subunits cause neonatal epilepsies including benign familial neonatal epilepsy (BFNE) and epileptic encephalopathy (EE) (rikee.org) ^14-17^. EE is a collection of epileptic syndromes accompanied by profound neurodevelopmental delay and psychomotor retardation ^18,19^.

K_v_7 channels are inhibited by membrane PIP_2_ depletion upon activation of G_q_-coupled receptors _2,6,13,20_. The underlying mechanism has been extensively investigated in K_v_7.1 ^3^. Voltage-clamp fluorometry studies have demonstrated that depolarization can activate the VSD of K_v_7.1 but fails to open the pore upon PIP_2_ depletion ^21^, suggesting that PIP_2_ is crucial for coupling the VSD to the pore domain. In the cryo-EM structure of K_v_7.1 channel in complex with KCNE3 and CaM, PIP_2_ interacts with the S2-S3 and S4-S5 linkers, and this interaction may facilitate the channel opening by straightening the unstructured loop between the S6 to Helix-A (pre-Helix-A) to a helix ^22^. In addition to the S2-S3 and S4-S5 linkers and S6 as potential PIP_2_ binding sites in K_v_7.1 ^21^, *in vitro* binding studies with helices A-D of K_v_7.1 have also identified basic residues in distal Helix-B that interact with PIP_2_ ^23^.

Despite the accumulating mechanistic insights into PIP_2_-dependent modulation of K_v_7.1 ^3^, it remains unclear whether neuronal K_v_7 channels are regulated by the same PIP_2_ binding residues and mechanism as K_v_7.1. There are several differences in PIP_2_-dependent modulation between K_v_7.1 and neuronal K_v_7 channels. First, PIP_2_ sensitivity of K_v_7.1 is regulated by KCNE1 ^24^, whereas PIP_2_ directly modulates neuronal K_v_7 channels without auxiliary β subunits ^13^. Second, previous electrophysiology studies with site-directed mutagenesis have suggested potential PIP_2_ binding sites unique to K_v_7.2 and K_v_7.3 including the regions between Helix-A and Helix-B (AB linker) and between Helix-B and Helix-C (BC linker) ^14,25-27^. Third, the AB linker of K_v_7.2 is much longer than that of K_v_7.1. Importantly, some epilepsy variants in K_v_7.2 and K_v_7.3 disrupt the channel sensitivity to the changes in cellular PIP_2_ level ^14,25,28,29^. Therefore, detailed investigation of how PIP_2_ regulates neuronal K_v_7 channels can increase our understanding of their physiological function in neurons and facilitate the development of new therapeutic strategies against epilepsy.

The first step toward understanding the molecular action of PIP_2_ on neuronal K_v_7 channels is to identify PIP_2_ binding sites. To achieve this, we employed all-atom molecular dynamics (MD) simulations. This technique has been successfully employed to provide atomic-level structural insights on lipid-protein interactions in membrane proteins with high spatiotemporal resolution ^30-34^, in close agreement with the experimental data ^30-35^. We chose to identify PIP_2_ binding sites in homomeric K_v_7.2 channels for several reasons. First, they produce robust K^+^ currents upon depolarization whereas homomeric K_v_7.3 channels are nonfunctional ^36-39^. Second, conditional deletion of K_v_7.2 but not K_v_7.3 results in cortical hyperexcitability, spontaneous seizures, and high mortality in mice ^40^. Third, there are significantly more epilepsy mutations found in *KCNQ2* than *KCNQ3* (ClinVar Database, NCBI) ^41,42^, and current suppression of homomeric K_v_7.2 channels is a common feature of EE variants of *KCNQ2* ^43^.

Here our MD simulations reveal multiple PIP_2_ binding sites in homomeric K_v_7.2 channels with more sites in the open state than the closed state. These sites include the S2-S3 linker, S4, S4-S5 linker, S6, pre-Helix-A, AB linker, Helix-B, and BC linker. Charge-neutralizing mutations of four PIP_2_-binding residues (R214Q in the S4, K219N in the S4-S5 linker, R325Q in pre-Helix-A, and R353Q in the AB linker) disrupt PIP_2_ binding to the mutated residues and decrease current densities of K_v_7.2 channels. Importantly, R214Q, K219N, and R325Q mutations reduced channel sensitivity to PIP_2_, with the triple R214Q/K219N/R353Q mutation inducing the largest effect. Our simulations further show that R214Q, K219N and R325Q mutations decouple the VSD activation from the pore domain of K_v_7.2 channels, while the R353Q mutation blocks the PIP_2_-induced bilayer-interacting conformation of helices A and B. These findings offer detailed mechanistic insights into PIP_2_-dependent modulation of K_v_7.2 channels.

## Results

### Differential PIP_2_ binding in closed and open K_v_7.2 channels

To identify PIP_2_ binding sites, we performed all-atom MD simulations on human K_v_7.2 channels in explicit lipid bilayers composed of phosphatidylcholine and PIP_2_. By adopting an integrative structural modeling approach using X-ray and cryo-EM data, we first constructed the open and closed states of K_v_7.2 channels within explicit lipid bilayers (Fig. 1a). The stability of the resulting models was investigated and reported in our previous publication ^14^. At the beginning of the lipid-binding simulations, 8 PIP_2_ lipid molecules (2.2% of the total lipid in each leaflet) were distributed around the channel with their starting positions randomized in each of the three independent, 500ns-long simulation replicates (Fig. 1b, Table 1). Differential binding of PIP_2_ lipids to open and closed channels was captured and presented as the PIP_2_ headgroup occupancy maps extracted from the entire simulation trajectory set for each state (Fig. 1c-d). Upon binding, PIP_2_ remained stably bound throughout the rest of the simulation time in all identified binding sites in both states (Fig. 1e-f).

**Table 1:**
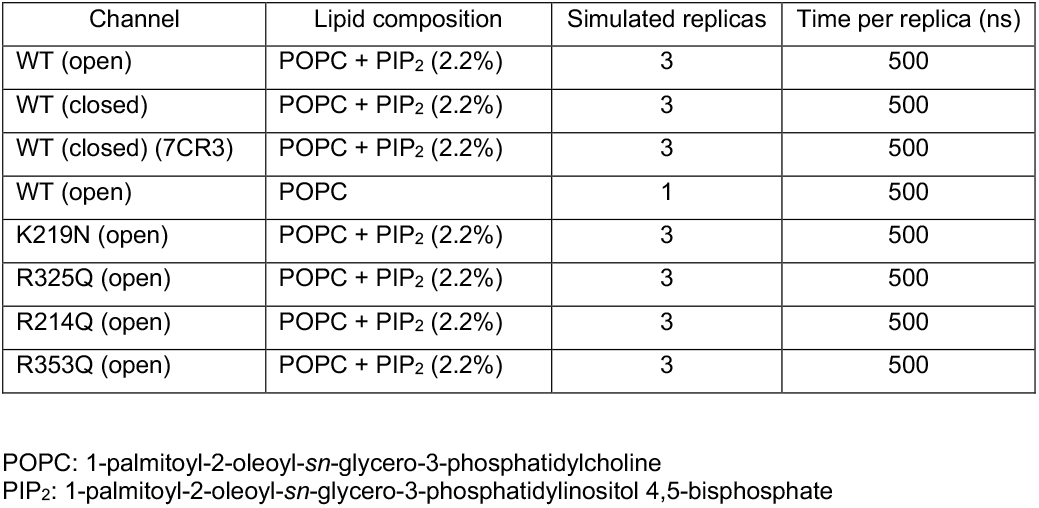
Details of the systems and the simulations performed.

**Figure 1.**
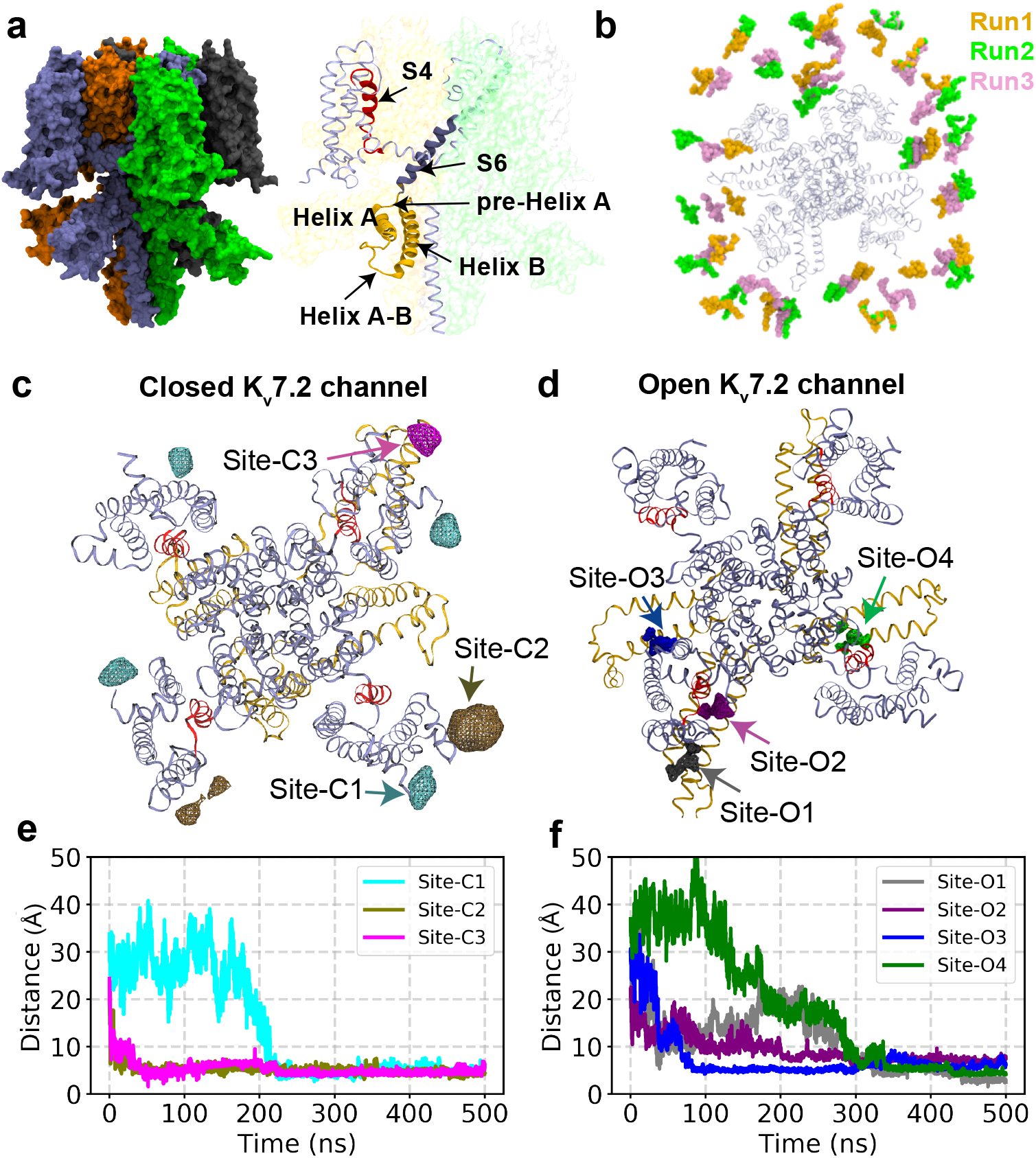
Simulation design for identifying PIP_2_ localization in open and closed K_v_7.2 channels. **(a)** Left: A tetrameric K_v_7.2 channel used in our simulations, where each subunit is highlighted with a different color in surface representation. Right: Detailed view of the important structural features. A tetrameric human K_v_7.2 channel in a closed state was modeled from the cryo-EM structure of human K_v_7.1 channel with a closed pore (PDB ID: 5VMS). For modeling the K_v_7.2 channel in an open state, TMD (targeted MD) was used to drive the transition of the closed K_v_7.2 state model towards the template based on the open state of K_v_1.2/ K_v_2.1 channel (PDB: 2R9R). **(b)** Top view of the simulation replicates (Runs) showing the initial placement of 8 PIP_2_ molecules in both upper and lower leaflets of the lipid bilayer surrounding K_v_7.2 channel in the absence of calmodulin. For each state/conformation (open or closed), 3 independent simulation replicates were performed in a tetrameric K_v_7.2 channel by randomly shuffling the initial positions of PIP_2_ lipids, as shown by different colors (Runs 1-3). **(c-d)** Volumetric maps of PIP_2_ headgroup occupancy extracted collectively from the simulation trajectories are shown as wireframes overlaid on the K_v_7.2 channel structure for the closed state **(c)** or the open state **(d)**. We label the PIP_2_ binding sites according to their proximity to the pore region. In the closed state, PIP_2_ headgroups bind to Site-C1 (N-terminal tail), Site-C2 (the S2-S3 linker), and Site-C3 (at the interface formed by Helix-A, the S2-S3 linker and the AB linker). In the open conformation, PIP_2_ headgroups are observed to bind Site-O1 (the interface formed by the N-terminal tail, the S2-S3 linker, distal Helix-B, and the BC linker), Site-O2 (distal Helix-B), Site-O3 (the interface formed by the S2-S3 linker, distal Helix-B, and the BC linker), and Site-O4 (the interface formed by the distal ends of the S4 and S6, the S4-S5 linker, and pre-Helix-A). **(e-f)** Distance between the center of mass (COM) of PIP_2_ headgroup and the COM of the binding site is plotted against simulation time. Initially all PIP_2_ molecules were placed at least 15 Å away from any of the binding sites captured in K_v_7.2 channel structure in the closed state **(e)** and the open state **(f)**. Once PIP_2_ lipids bind to any of the binding sites, they remained stably bound throughout the simulation time.

We discover that the PIP_2_ headgroup interacts with 3 distinct sites in the closed state (Fig. 1c): Site-C1 (intracellular N-terminal tail), Site-C2 (the S2-S3 linker), and Site-C3 (the interface formed by Helix-A, the S2-S3 and AB linkers) (Fig. 1c). Notably, repeating the lipid-binding simulations on the recently solved cryo-EM structure of a closed K_v_7.2 channel ^44^ verified the formation of the same PIP_2_ binding sites (Supplementary Fig. S1). In the open state, we have identified four distinct PIP_2_-binding sites, all enriched with basic residues: Site-O1 (the interface formed by the S2-S3 and AB linkers), Site-O2 (distal Helix-B), Site-O3 (the interface formed by the N-terminal tail, the S2-S3 linker, distal Helix-B, and the BC linker), and Site-O4 (the interface formed by the ends of S4 and S6, the S4-S5 linker, and pre-Helix-A) (Fig. 1d).

Although PIP_2_ interact with the N-terminal tail and the S2-S3 and AB linkers in both closed and open states, there are key differences in the PIP_2_ binding sites of these two states. PIP_2_ headgroup occupancies at the four sites in the open channel are more focused than those at the three sites in the closed channel (Fig. 1c-d). Importantly, PIP_2_ localizes to the periphery of the VSD in the closed channel, whereas PIP_2_ binding spreads to a larger area in the open channel including the cytoplasmic ends of the VSD and S6, pre-Helix-A, distal Helix-B, and the BC linker (Fig. 1c-d), suggesting that the opening of K_v_7.2 channels involves PIP_2_ interaction with VSD, the gate, and intracellular helices.

### Characterizing specific lipid-protein interactions in K_v_7.2 channels

To map PIP_2_-binding sites, we analyzed lipid–protein interactions during the last 200ns of MD trajectories by quantifying the contact probability between each moiety of the PIP_2_ headgroup and the PIP_2_- binding residues. In the closed channel, Site-C1 is formed by a cluster of basic residues (K76, R87, and R89) in the intracellular N-terminal tail. R87 and R89 exclusively interact with P5-phosphate, whereas K76 establishes contacts with all the hydroxyl and phosphate groups on the inositol ring in PIP_2_ (Supplementary Fig. S2a, d). In Site-C2, a cluster of basic residues in the S2-S3 linker (R153, R158, and K162) coordinate PIP_2_. R153 and K162 show high contact probabilities to P4-phosphate and P5-phosphate, whereas R158 interacts with all groups in PIP_2_ (Supplementary Fig. S2b, e). Site-C3 is formed by R155 in the S2-S3 linker, Y347 in Helix-A, and R353 in the AB linker. R155 has high contact probabilities for P5-phosphate and the hydroxyl group at position 6 of the inositol ring, whereas R353 interacts preferentially with P4- and P5-phosphates (Supplementary Fig. S2c, f).

In the open channel, Site-O1 is formed by residues in the S2-S3 linker (K162 and R165) and the AB linker (F346, Y347, and R353) (Fig. 2a, b). All the three basic residues in this site show high contact probabilities for P5-phosphate on PIP_2_ headgroup (Fig. 2a). In Site-O2, a cluster of basic residues in distal Helix-B (K552, R553, and K554) coordinate PIP_2_ (Fig. 2b, f). K552 and R553 primarily bind to P5-phosphate, while K554 interacts with P4- and P5-phosphates and hydroxyl group at position 3 of the inositol ring (Fig. 2b). Site-O3 is formed by residues in the intracellular N-terminal tail (R87), the S2-S3 linker (R153, Y154, and K166), distal Helix-B (R553), and the BC linker (R560) (Fig. 2c, g). Y154 and K166 predominantly interacts with P4-phosphate, whereas the other basic residues bind to P5-phosphate. In Site-O4, PIP_2_ interacts with R214 at the end of S4, K219 in the S4-S5 linker, K319 and Q323 in distal end of S6, and R325 in pre-Helix-A (Fig. 2d, h). Most of these residues, except K319, show high contact probabilities for P5-phosphate in PIP_2_.

**Figure 2.**
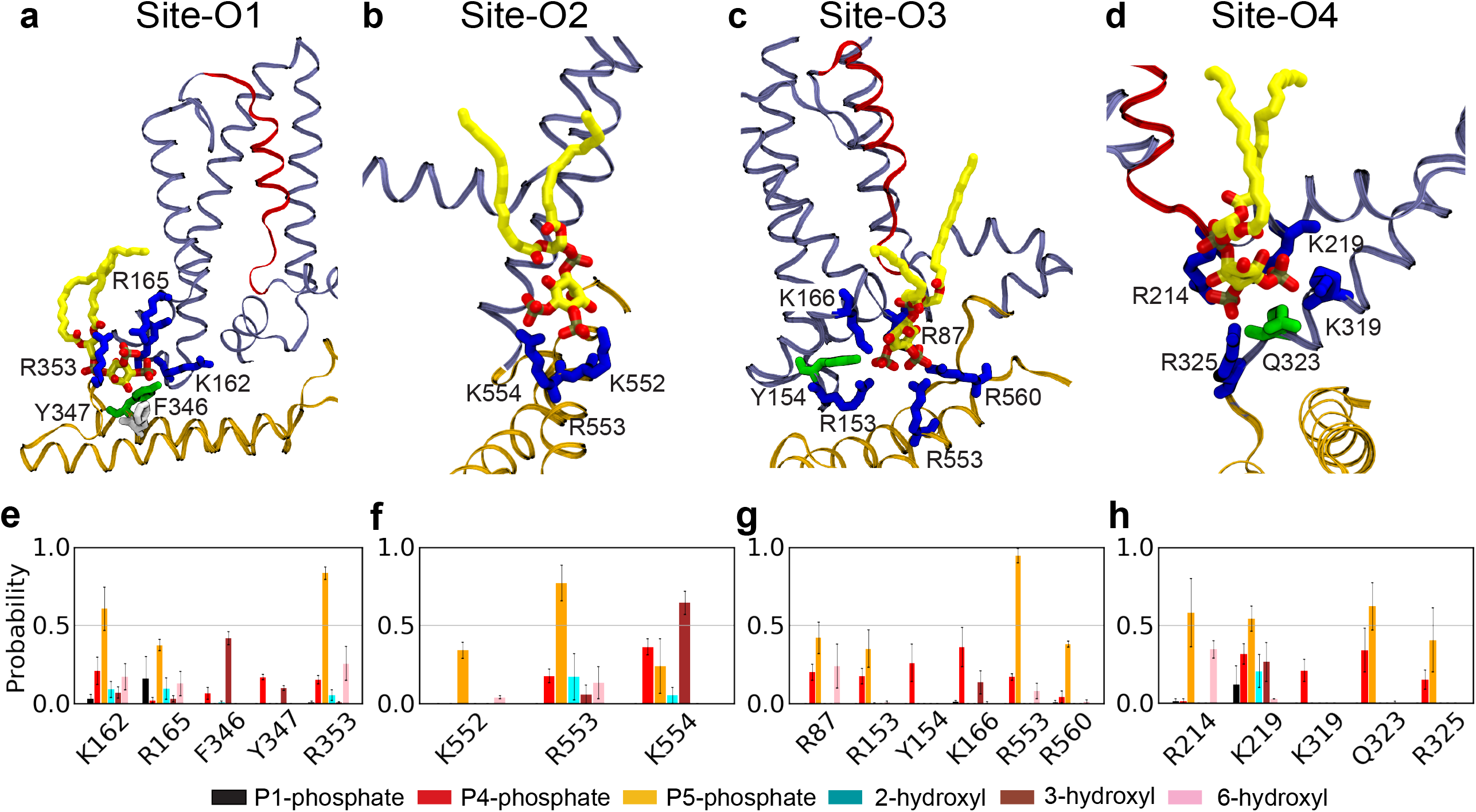
PIP_2_ coordination in 4 binding sites captured in the modeled structure of the open K_v_7.2 channel. **(a-d)** PIP_2_ coordination in Site-O1 **(a)**, Site-O2 **(b)**, Site-O3 **(c)**, and Site-O4 **(d)**. K_v_7.2 protein is shown in ribbon representation with S4 in red, helices A and B in brown, and the rest of the protein in ice blue. A PIP_2_ lipid (carbon atoms in yellow, oxygen in red, and phosphorus atoms in tan) and the residues of the binding pocket at each site at the end of simulations are shown in sticks (basic residues in blue, polar in green, acidic in red, and hydrophobic in white). **(e-h)** The contact probability for each chemical moiety of PIP_2_ headgroup with the key residues in the binding Site-O1 **(e)**, Site-O2 **(f)**, Site-O3 **(g)**, and Site-O4 **(h)**. The chemical moieties of PIP_2_ headgroup include P1-phosphate (black), P4-phosphate (red), P5-phosphate (orange), 2-hydroxyl (cyan), 3-hydroxyl (brown), and 6-hydroxyl (pink) group of the inositol ring. A heavy-atom distance cutoff of 4 Å was used to define a contact between a protein residue and a phosphate group of PIP_2_, whereas a 3.5 Å cutoff was used to define a contact between a residue and a hydroxyl group of PIP_2_. Analysis of the contact probabilities was performed over the last 200 ns of the simulation trajectories. Data represent mean ± SEM for 3 independent trajectories.

Overall, PIP_2_ binds to many more basic residues in the open state than the closed state. However, both states share 4 common PIP_2_ binding residues including R87 at the N-terminal tail, R153 and K162 in the S2-S3 linker, and R353 in the AB linker. Interestingly, the simulation trajectories (Supplementary Movies 1-3) show that R353 comes in contact with PIP_2_ first after which the lipid forms contacts with other residues in Site-O1, suggesting that R353 might act as an initial anchor point of PIP_2_ in Site-O1.

### Charge-neutralizing mutations of potential PIP_2_ binding residues disrupt voltage-dependent activation of K_v_7.2 channels

PIP_2_ is required for activation of all K_v_7 channels^3^. To test the functional impact of PIP_2_ binding sites identified by our MD simulations, we introduced charge-neutralizing mutations in select residues that had high contact probability to PIP_2_ headgroups. In Site-O4, we introduced R214Q in the distal S4, K219N in the S4-S5 linker, and R325Q in pre-Helix-A (Fig. 3a-i). Since MD simulations identified R353 as an initial anchoring point for PIP_2_ in Site-O1, we also made R353Q in the AB linker (Fig. 3j-l). To test if charge-neutralizing mutations disrupt PIP_2_ binding to mutated basic residues, we performed 500-ns MD simulations after introducing each mutation in the PIP_2_-bound conformation of wild-type (WT) channels and monitored the distance between the PIP_2_ headgroup and the mutated residues.

**Figure 3.**
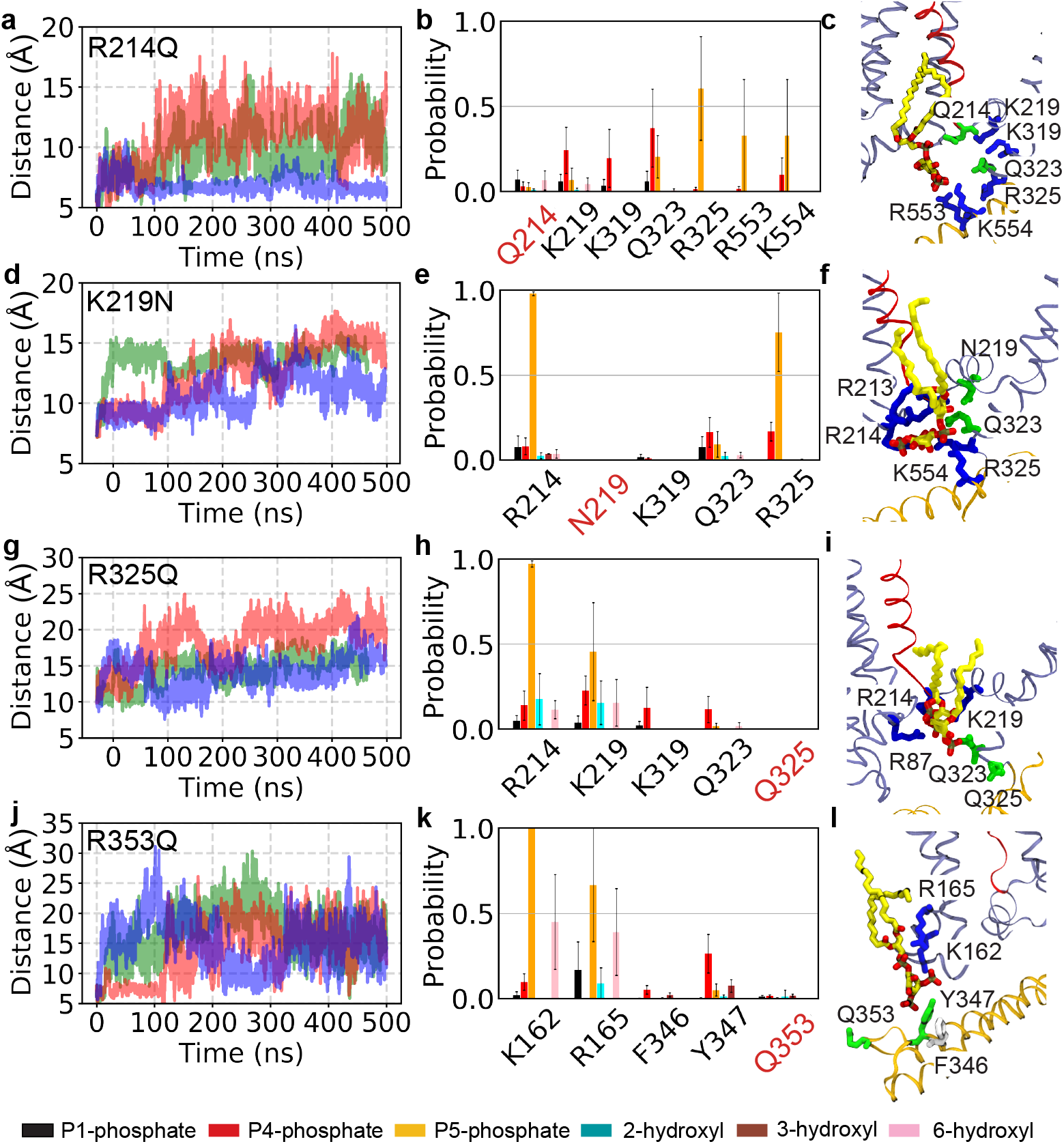
Charge-neutralizing mutations disrupt PIP_2_ binding to the mutated residues in the open K_v_7.2 channel. **(a, d, g, j)** Time evolution of the distance between the COM of PIP_2_ headgroup and the COM of the mutated residue upon introducing mutations including R214Q **(a)**, K219N **(d)**, R325Q **(g)**, and R353Q **(j)** in PIP_2_-bound K_v_7.2 channel in an open state from three independent simulations (different colors). Introduction of these mutations resulted in the dissociation of bound PIP_2_ lipid, as indicated by the distance increase. **(b, e, h, k)** The contact probability for each chemical moiety of PIP_2_ headgroup with the key residues in open K_v_7.2 channels containing R214Q **(b)**, K219N **(e)**, R325Q **(h)**, and R353Q **(k)**. The chemical moieties of PIP_2_ headgroup include P1-phosphate (black), P4-phosphate (red), P5-phosphate (orange), 2-hydroxyl (cyan), 3-hydroxyl (brown), and 6-hydroxyl (pink) group of the inositol ring. A heavy-atom distance cutoff of 4 Å was used to define a contact between a residue and a phosphate group of PIP_2_, whereas a 3.5 Å cutoff was used to define a contact between a residue and a hydroxyl group of PIP_2_. Analysis of the contact probabilities was performed over the last 200 ns of the simulation trajectories. Data represent mean ± SEM for 3 independent trajectories. **(c, f, i, l)** PIP_2_ coordination in Site-O4 of K_v_7.2-R214Q **(c)**, K_v_7.2-K219N **(f)**, and K_v_7.2-R325Q **(i)** and in Site-O1 of K_v_7.2-R353Q (**l**) channel at the end of the respective simulations. The protein is shown in ribbon representation with the S4 in red, helices A and B in brown, and the rest of the protein in ice blue. A PIP_2_ lipid (carbon atoms in yellow, oxygen in red, and phosphorus in tan) and the residues of the binding pocket at each site are shown in sticks (basic residues in blue, polar in green, acidic in red, and hydrophobic in white).

Upon introducing R214Q, we observed that the bound PIP_2_ dissociated from the mutated residue and Site-O4 and diffused to basic residues in distal Helix-B in 2 out of 3 simulations (Fig. 3a-c). The introduction of K219N or R325Q resulted in dissociation of PIP_2_ from the mutated residue in all simulations (Fig. 3d-i). Compared to the WT channel (Fig. 2d, h), these mutations also decreased PIP_2_ binding to K319 and Q323 (Fig. 3e-f, h-i) but not to other basic residues in Site-O4 (Fig. 3e-f, h-i). Introduction of R353Q resulted in dissociation of PIP_2_ from the mutated residue in all simulations, although PIP_2_ remained bound to K162 and R165 in Site-O1 (Fig. 3j-l).

To test if these mutations affect voltage-gated activation of K_v_7.2 channels, we performed whole cell patch clamp recording in CHO hm1 cells ^14,25,45^, which display depolarized resting membrane potential (V_m_) and reversal potential (E_rev_) due to a low level of endogenous K^+^ channels ^45,46^. Application of depolarizing voltage steps from -100 to +20 mV in cells transfected with GFP and WT K_v_7.2 produced a slowly activating outward K^+^ current with peak current density of 26.5 ± 1.4 pA/pF at +20 mV (Fig. 4a-c, Supplementary Fig. S3). Current activation was sigmoidal, and full activation was reached from 0 mV step with half-maximal current activation potential (V_1/2_) of -30.5 ± 0.5 mV (Fig. 4a-d, Table 2). Due to this increase in outward K^+^ currents, these cells also displayed hyperpolarized V_m_ and E_rev_ compared to cells transfected with GFP alone (Supplementary Table S1).

**Table 2.** Biophysical properties of K_v_7.2 homomers in CHO hm1 cells co-transfected with GFP and PIP5K.

| Transfection | *n* | *Leak subtracted I*<br>at +20 mV (pA) | $G/G_{\max}$<br>$V_{1/2}$<br>(mV) | $G/G_{\max}$<br>*k*<br>(mV/e fold) |
| --- | --- | --- | --- | --- |
| GFP | 12 | 27.3 ± 4.0* | -65.8 ± 9.2* | 0.7 ± 0.2* |
| GFP + WT | 31 | 567.7 ± 43.3 | -30.5 ± 0.5 | 5.9 ± 0.7 |
| GFP + R214Q | 16 | 299.6 ± 35.5* | -31.4 ± 0.5 | 4.3 ± 0.4* |
| GFP + K219N | 12 | 150.4 ± 32.4* | -13.7 ± 1.7* | 6.5 ± 0.7 |
| GFP + R325Q | 11 | 63.9 ± 7.5* | -65.1 ± 3.6* | 1.2 ± 0.4* |
| GFP + R353Q | 13 | 310.2 ± 39.9* | -31.7 ± 1.1 | 5.0 ± 1.1 |
| GFP + R214Q/K219N | 15 | 284.1 ± 19.5* | -9.7 ± 0.6* | 5.6 ± 0.4* |
| GFP + R214Q/K219N/R353Q | 14 | 312.4 ± 38.5* | -9.8 ± 1.0* | 5.5 ± 0.6 |
| GFP + WT + PIP5K | 33 | 2164.0 ± 92.4<sup>^</sup> | -40.3 ± 0.6<sup>^</sup> | 2.3 ± 0.3<sup>^</sup> |
| GFP + R214Q + PIP5K | 15 | 1500.8 ± 138.3<sup>^†</sup> | -38.7 ± 0.7<sup>^</sup> | 2.2 ± 0.4<sup>^</sup> |
| GFP + K219N + PIP5K | 12 | 981.9 ± 92.9<sup>^†</sup> | -29.4 ± 0.8<sup>^†</sup> | 3.2 ± 0.6<sup>^</sup> |
| GFP + R325Q + PIP5K | 11 | 75.0 ± 9.2<sup>†</sup> | -33.3 ± 3.0<sup>^†</sup> | 8.5 ± 1.4<sup>^†</sup> |
| GFP + R353Q + PIP5K | 16 | 1373.5 ± 79.5<sup>^†</sup> | -42.4 ± 1.2<sup>^</sup> | 1.2 ± 0.3<sup>^</sup> |
| GFP+ R214Q/K219N+ PIP5K | 19 | 997.7 ± 70.4<sup>^†</sup> | -20.5 ± 0.9<sup>^†</sup> | 3.8 ± 0.6<sup>^†</sup> |
| GFP+ R214Q/K219N/R353Q + PIP5K | 14 | 492.2 ± 48.4<sup>†</sup> | -20.8 ± 1.2<sup>^†</sup> | 4.3 ± 0.9<sup>^</sup> |

*n*, number; NA, not applicable; *Leak subtracted peak current* measured at +20 mV; $V_{1/2}$ , half-activation potential; *k*, the slope factor; $\tau$ , activation time constant measured at +20 mV. All values are calculated from leak subtracted current. $V_{1/2}$ and *k* are calculated from normalized conductance $G/G_{\max}$ . Mean ± SEM (GFP+ selected variant: \**p*<0.05 for K<sub>v</sub>7.2 WT vs. mutant; <sup>†</sup>*p*<0.05 for K<sub>v</sub>7.2 WT + PIP5K vs. mutants + PIP5K; <sup>^</sup>*p*<0.05 for the difference between -PIP5K and +PIP5K within the same transfection.

**Figure 4.**
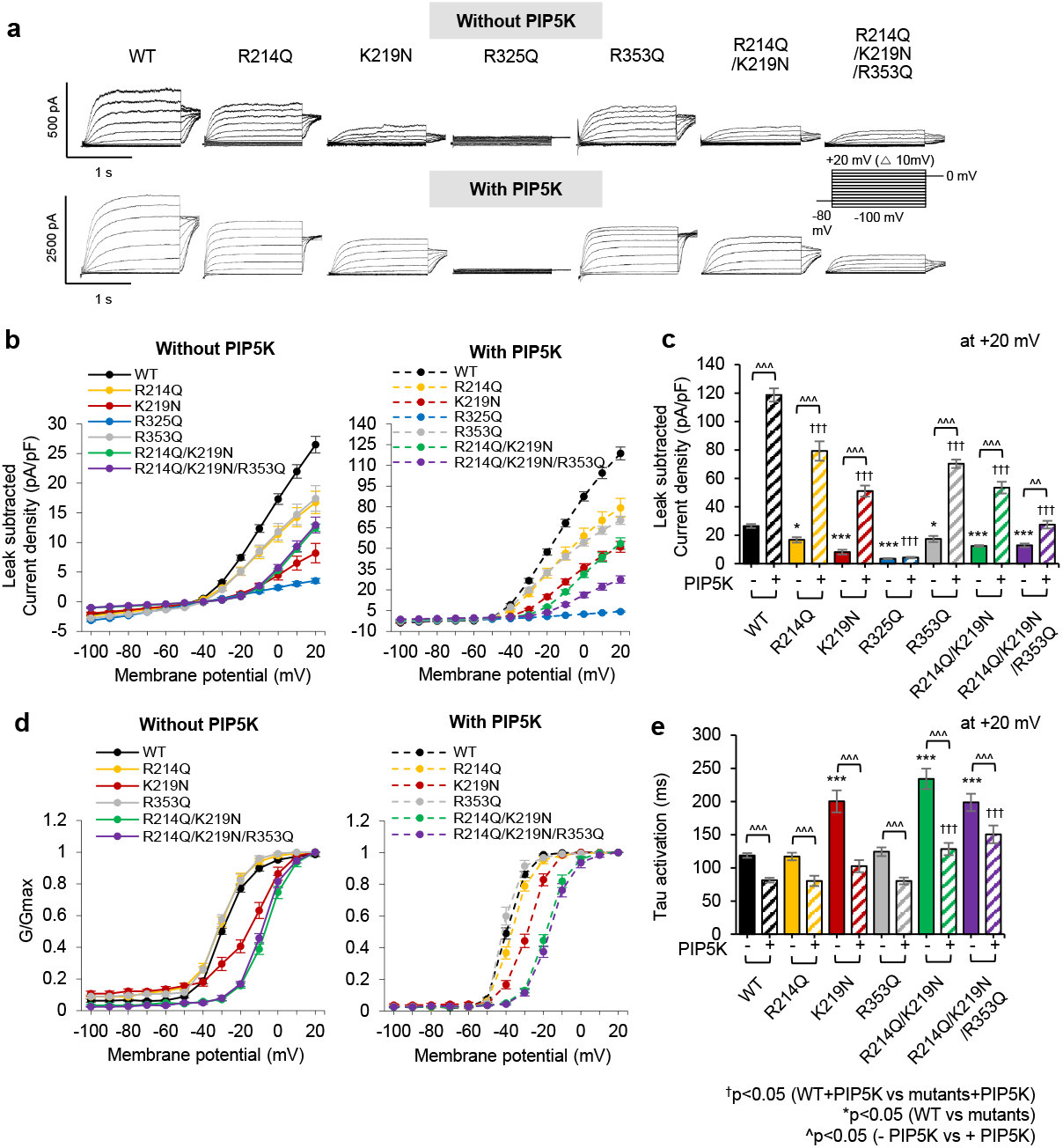
Charge-neutralizing mutations of potential PIP_2_ binding residues decrease voltage- and PIP_2_-dependent activation of homomeric K_v_7.2 channels. Whole cell voltage clamp recordings of macroscopic K^+^ currents through K_v_7.2 WT or mutant channels were performed in CHO hm1 cells cotransfected with GFP and PIP5K. The DNA plasmid ratio was 0.2:0.45:0.45 (GFP: K_v_7.2 WT or mutant: PIP5K). Cells were held at -80 mV. K^+^ currents were evoked by depolarizing voltage steps for 1.5 s from -100 mV to +20 mV in 10-mV increments, followed by a step to 0 mV for 300 ms. **(a)** Representative traces after subtraction of leak currents. Leak current was defined as non-voltage-dependent current from GFP-transfected cells. Note different Y-axis scales for systems with and without PIP5K. **(b)** Average peak current densities (pA/pF) of K_v_7.2 WT or mutant channels with or without PIP5K coexpression at all voltage steps. **(c)** Average peak current densities of WT or mutant K_v_7.2 channels at +20 mV. p values are computed from one-way ANOVA post-hoc Fisher’s test. **(d)** Normalized conductance (G/G_max_) at all voltage steps. **(e)** Average activation constant (*τ*) at +20 mV. The number of GFP-positive cells that were recorded without PIP5K coexpression: K_v_7.2 WT (n=31), R214Q (n=16), K219N (n=12), R325Q (n=11), R353Q (n=13), R214Q/ K219N (n=15), R214Q/K219N/R353Q (n=14). The number of GFP-cotransfected cells that were recorded with PIP5K: K_v_7.2 WT (n=33), R214Q (n=15), K219N (n=12), R325Q (n=11), R353Q (n=16), R214Q/ K219N (n=19) or R214Q/K219N/R353Q (n=14). Data represent the mean ± SEM. One-way ANOVA with post-hoc Fisher’s multiple comparison test was used. GFP+ selected variant: *p<0.05 for K_v_7.2 WT vs. mutant (**p<0.01, ***p<0.005); ^†^p<0.05 for K_v_7.2 WT + PIP5K vs. mutants + PIP5K (^††^p<0.01, ^†††^p<0.005); ^ p<0.05 for the difference between -PIP5K and +PIP5K within the same transfection (^ ^ p<0.01, ^ ^ ^ p<0.005).

Compared to WT channels, K_v_7.2-K219N channels produced K^+^ currents with significantly smaller peak current density (8.2 ± 1.6 pA/pF at +20 mV), a large depolarizing shift in V_1/2_ (−13.7 ± 1.7 mV), and a slower activation kinetic (Fig. 4a-e, Table 2). The R325Q mutation significantly reduced peak current density by ∼90% (3.5 ± 0.4 pA/pF at +20 mV) (Fig. 4a-c). The peak current densities of K_v_7.2-R214Q and K_v_7.2-R353Q channels were also decreased by ∼30%, but their voltage dependence and activation time constant were unaffected (Fig. 4a-e, Supplementary Fig. S3). Similar surface expressions were observed for WT and all tested mutants except for K_v_7.2-R214Q (Fig. 5), which displayed lower surface expression than the WT (61.8 ± 11.0% of WT, Fig. 5a, c). The surface/total protein ratio of all tested mutants did not differ from WT K_v_7.2 (Fig. 5e), suggesting that none of the mutations affected the proportion of K_v_7.2 to express on the plasma membrane.

**Figure 5.**
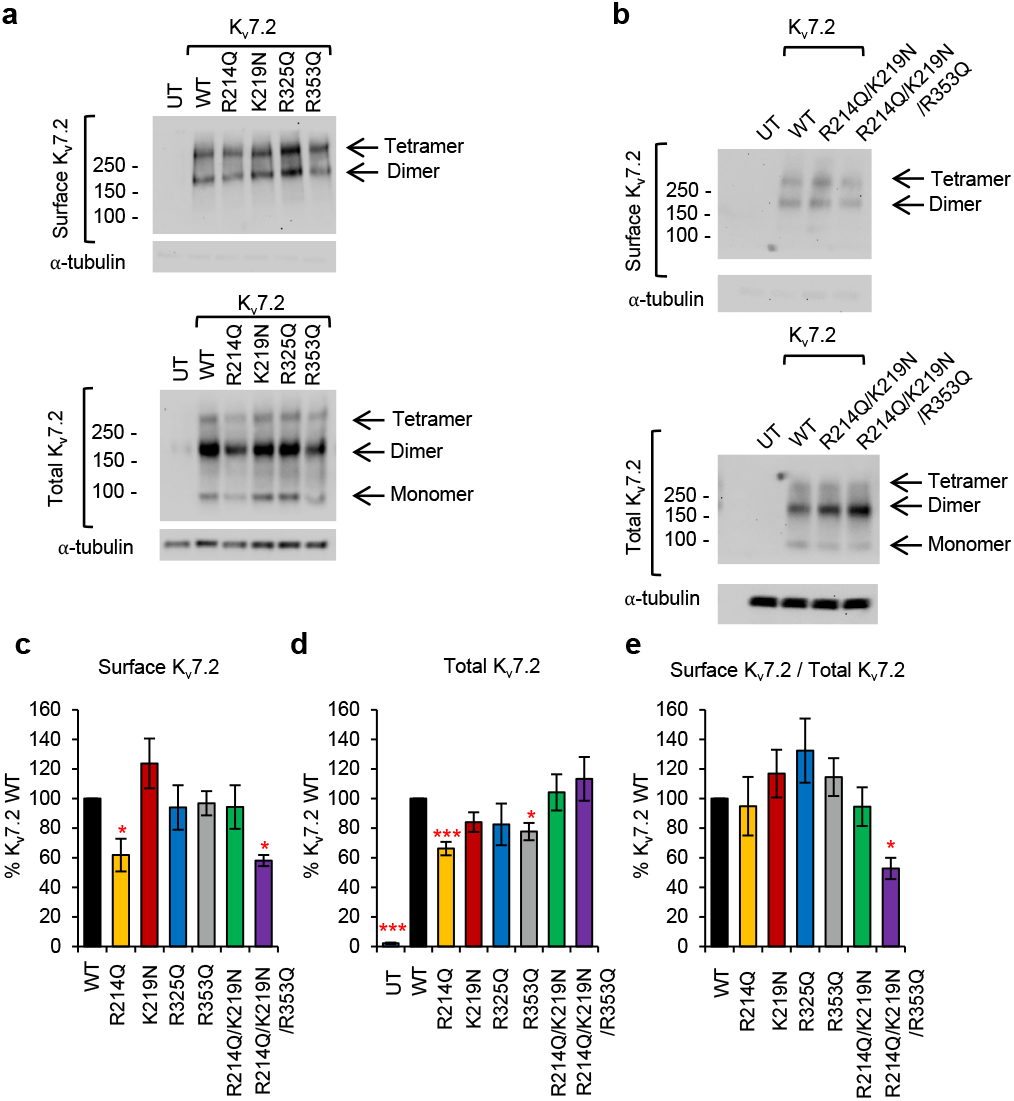
Charge-neutralizing mutations of potential PIP_2_ binding residues are expressed on the cell surface of CHO hm1 cells. Surface biotinylation was performed on live CHO hm1 cells with Sulfo-NHS-SS-Biotin. Surface biotinylated proteins were isolated by NeutrAvidin agarose beads, released from the beads by a 30-min incubation in the Sample Buffer at 75 °C, and analyzed by immunoblot analysis. **(a-b)** Representative immunoblot images of surface and total K_v_7.2 WT and mutant proteins. Untransfected cells (UT) served as a negative control. In total lysate (bottom blot), WT and mutant K_v_7.2 proteins were expressed as monomers, dimers, and tetramers as we have previously reported ^14^. In surface biotinylated fractions (top blots), K_v_7.2 proteins were detected only as dimers and tetramers. The lack of monomeric bands is due to the extended exposure to the denaturing Sample Buffer, which facilitates oligomerization of K_v_7.2 proteins. **(c)** Quantification of surface K_v_7.2, which was normalized to total *α*-tubulin and presented as % K_v_7.2 WT (n=4 for each construct). **(d)** Total K_v_7.2 level was normalized to the loading controls (GAPDH, *α*-tubulin, or *β*-tubulin). The numbers are (n=9) for single mutations, (n=4) for the double-mutant R214Q/K219N, and (n=4) for the triple-mutant R214Q/K219N/R353Q. Full blots are illustrated in Supplementary Fig. S5. **(e)** Quantification of the “surface K_v_7.2 / total K_v_7.2” ratio, which was normalized to the WT ratio (WT as 100%, n=4 for each construct). Data represent the mean ± SEM. One-way ANOVA with post-hoc Fisher’s test was conducted. *p<0.05, **p<0.01, ***p<0.005.

### Charge-neutralizing mutations of potential PIP_2_ binding residues alter PIP_2_ sensitivity of K_v_7.2 channels

To test if charge-neutralizing mutations alter gating modulation of K_v_7 channels by PIP_2_, we increased cellular PIP_2_ level by co-transfecting phosphatidylinositol-4-phosphate 5-kinase (PIP5K) which converts phosphatidylinositol 4-phosphate to PIP_2 47_. Since the endogenous membrane level of PIP_2_ is not enough to saturate K_v_7 channel activation ^24^, enhancing cellular PIP_2_ level by PIP5K expression is shown to increase single-channel open probability ^48^ and whole-cell current densities of K_v_7.2 channels ^14,28^. Consistent with previous reports ^14,28,48^, PIP5K expression significantly increased peak K_v_7.2 current density by 4.4 ± 0.2-fold (118.7 ± 4.7 pA/pF) with a hyperpolarizing shift in their voltage dependence and a faster activation kinetic (Fig. 4a-e, Supplementary Figs. S3-S4).

In contrast, the R325Q mutation abolished the PIP5K-induced current potentiation (Fig. 4a-d, Supplementary Figs. S3-S4). In the presence of PIP5K, K_v_7.2 channels containing R214Q, K219N or R353Q mutations produced significantly less outward K^+^ currents compared to WT channels (K_v_7.2-R214Q: 79.3 ± 6.8 pA/pF; K_v_7.2-K219N: 51.1 ± 3.9 pA/pF; K_v_7.2-R353Q: 70.3 ± 3.0 pA/pF) (Fig. 4a-e, Supplementary Figs. S3-S4, Table 2). However, the fold increases in their PIP5K-induced current potentiation were comparable to the WT channels (4.7 ± 0.4 fold for R214Q, 6.2 ± 0.5 fold for K219N, 4.0 ± 0.2 fold for R353Q) (Fig. 4a-e, Supplementary Figs. S3-S4, Table 2), suggesting that the charge-neutralizing mutation of a single residue may not be sufficient to fully dissociate PIP_2_ from Site-O4 and Site-O1 (Fig. 3b,e,h,k).

Therefore, we next generated the R214Q/K219N double mutant (DM) and the R214Q/K219N/R353Q triple mutant (TM). Compared to K_v_7.2-WT channels, both DM and TM channels displayed smaller peak current densities (DM = 12.5 ± 0.6 pA/pF, TM = 12.9 ± 1.3 pA/pF) and activated at more depolarized voltages with slower activation kinetics (Fig. 4a-e, Supplementary Figs. S3-S4, Table 2). PIP5K co-expression increased the peak current density of DM channels by 4.3 ± 0.3-fold (53.5 ± 4.1 pA/pF) but that of TM channels only by 2.1 ± 0.2-fold (27.5 ± 2.7 pA/pF) (Fig. 4a-e, Supplementary Figs. S3-S4, Table 2), indicating that the triple mutation decreased the channel sensitivity to PIP_2_ enhancement.

To further investigate PIP_2_ sensitivity of mutant channels, we examined current decay upon membrane PIP_2_ depletion. To achieve this, we coexpressed *Danio rerio* voltage-sensitive phosphatase (Dr-VSP) ^14^. Upon activation of Dr-VSP, K^+^ currents through WT channels reached a maximal decay of at +100 mV (0.52 ± 0.02, Fig. 6a-c). K_v_7.2-R353Q channels displayed larger current decays at +40 and +60 mV, but showed similar decays to WT channels at +100 mV (Fig. 6a-c). In contrast, the VSP-induced current decays of R214Q and K219N channels were smaller than WT channels and were minimal in K_v_7.2-R325Q, DM, and TM channels (Fig. 6a-c), indicating these mutants have decreased sensitivity to PIP_2_ depletion.

**Figure 6.**
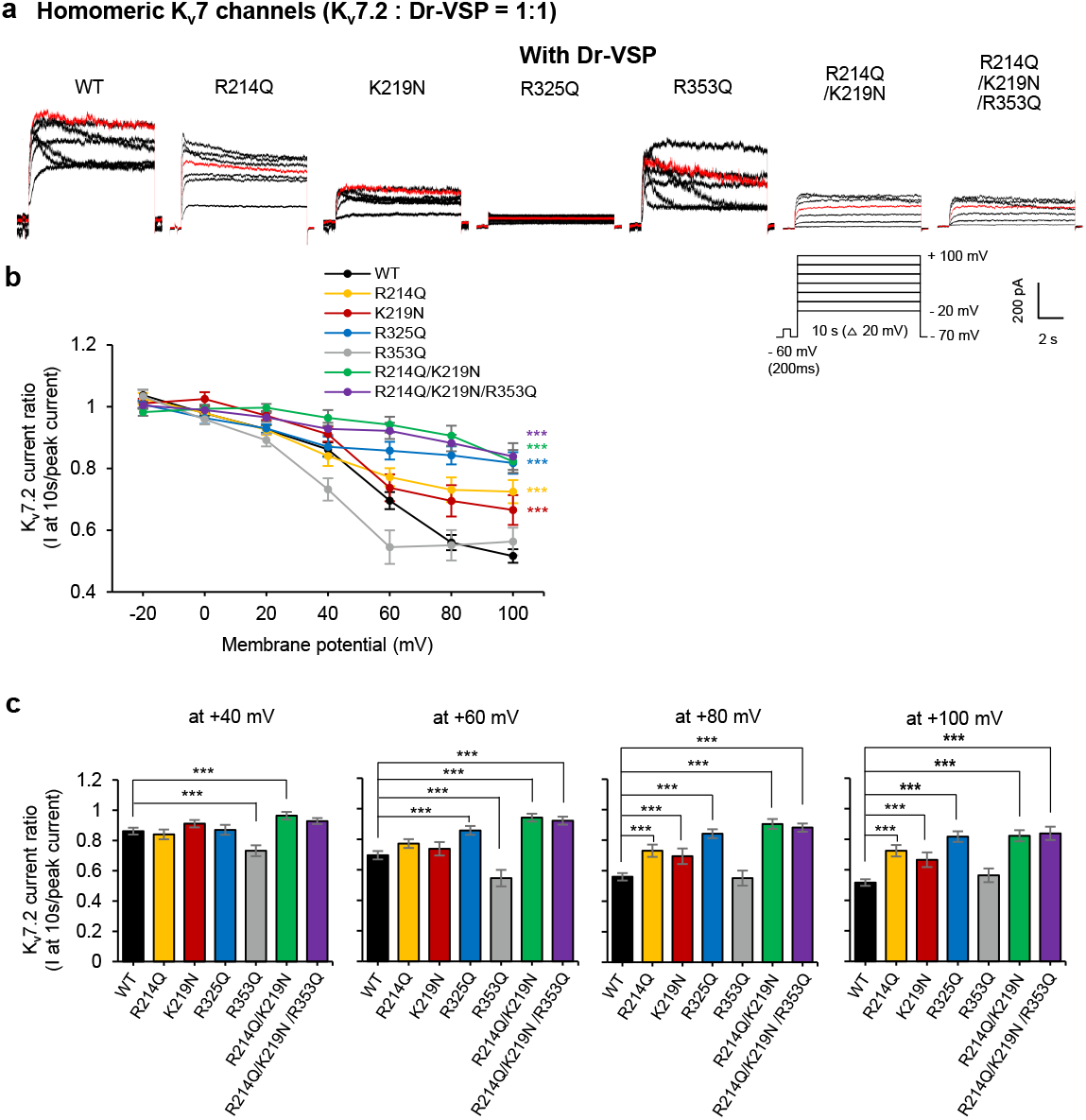
Charge-neutralizing mutations of potential PIP_2_ binding residues alter K_v_7.2 current response to Dr-VSP activation. **(a)** Representative current traces showing Dr-VSP-mediated K_v_7.2 current decay in CHO hm1 cells co-expressing Dr-VSP and K_v_7.2 WT or mutants from -20 mV to +100 mV. CHO cells were held at –70 mV, and a brief voltage step to –60 mV was applied to calculate the linear leak. 10 s step depolarizations were applied in 20-mV steps from –20 to +100 mV with 2 min inter-step intervals to allow PIP_2_ regeneration. Red trace shows current decay curve when cells were held at +40 mV. **(b)** Ratio of current decay in K_v_7.2 WT or mutant channels at each voltage step. **(c)** K_v_7.2 current decay ratio at +40 mV, +60 mV, +80 mV and +100 mV. The number of cotransfected cells that were recorded with EGFP-tagged Dr-VSP: K_v_7.2 WT (n=27), R214Q (n=14), K219N (n=13), R325Q (n=12), R353Q (n=15), R214Q/ K219N (n=13) or R214Q/K219N/R353Q (n=11). Data represent the mean ± SEM. One-way ANOVA Fisher’s test results are shown (*p <0.05, **p < 0.01 and ***p < 0.005).

### PIP_2_-mediated correlated motions of the VSD and the pore domain

Voltage-dependent conformational changes of the VSD and the pore domain are critical for the gating of K_v_ channels ^49^. To examine PIP_2_-mediated allosteric interactions, we investigated the communities formed in the WT and mutant channels by employing dynamic network analysis and Pearson correlation. These communities correspond to sets of residues that move in a correlated manner during the MD simulations and thus represent strongly connected regions. We detected the presence of a pronounced community connecting the end of S4, the S4-S5 linker, S6, and pre-Helix-A in WT channels (Fig. 7a). The thickness of the edges connecting the amino acids corresponds to the strength of the correlation between them. All mutant channels show thinner edges within this community (Fig. 7b-d), indicating weaker correlated motions in this area compared to WT channels. Interestingly, we also detected the emergence of additional (sub)communities in all the studied mutants (Fig. 7b, c, d), in line with the described weaker interactions in this region. The presence of disconnected communities suggests uncorrelated dynamics of this region (Site-O4) after decreased PIP_2_ binding (Fig. 3). These data suggest that R214Q, K219N, and R325Q mutations disrupt the correlated motions of the VSD and the S6 gate in the pore domain of K_v_7.2.

**Figure 7.**
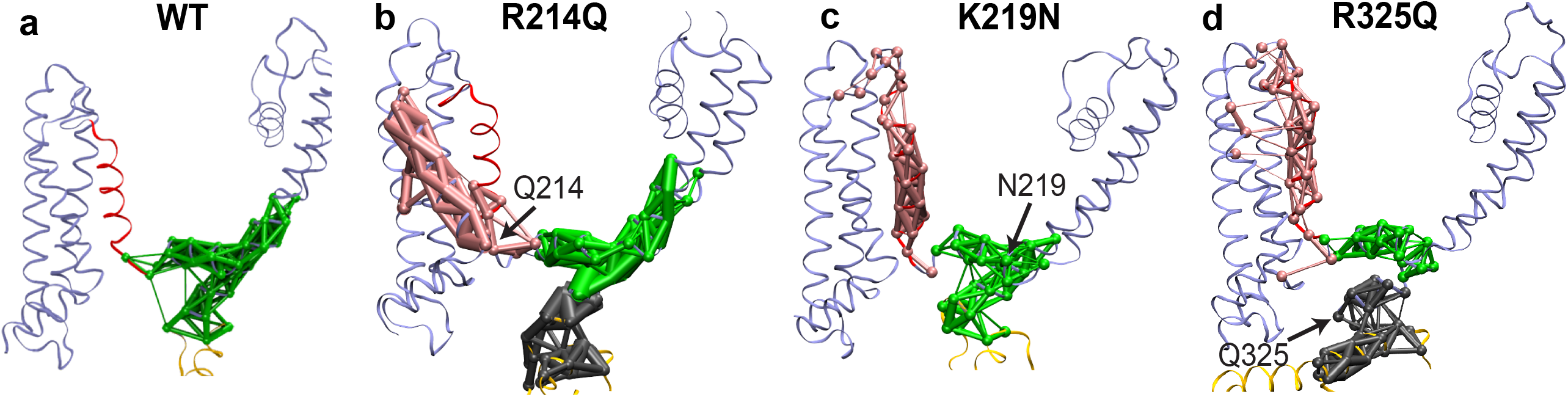
Charge-neutralizing mutations of potential PIP_2_ binding residues in Site-O4 disrupt the correlated motions of the VSD and the S6 gate of K_v_7.2 channels. Network-based community analysis in WT **(a)** and mutant K_v_7.2 channels containing R214Q **(b)**, K219N **(c)**, and R325Q **(d)** mutations in Site-O4. A community represents a set of residues that move in a correlated manner during the MD simulations. The thickness of the edges between the amino acids within each community corresponds to the strength of correlation between them. The network analysis was performed after combining all 3 independent simulation trajectories. **(a)** Binding of PIP_2_ at Site-O4 in the WT channel results in correlated motions of the VSD (the S4 and the S4-S5 linker) and the gate S6, highlighted by a single community in green. **(b)** Introduction of the R214Q mutation leads to uncorrelated motions of the VSD and the gate, highlighted by 3 separate communities colored in pink, green, and grey. **(c)** Introduction of the K219N mutation leads to uncorrelated motions of the VSD and the gate, highlighted by the smaller community in green and the emergence of an additional community colored in pink. **(d)** K_v_7.2-R325Q channels displayed weaker correlated motions of the S4 and the S4-S5 linker, and the decoupling of the S6 of the pore domain from the VSD, highlighted by the much smaller community in green and the emergence of 2 additional communities colored in pink and grey.

### PIP_2_-dependent conformational change of helices A and B

The cryo-EM structure of K_v_7.1/KCNE3 in complex with CaM suggests that PIP_2_ induces a conformational change in the cytoplasmic domain that may facilitate the opening of K_v_7.1 channels ^22^. To test the conformational impact of PIP_2_ on K_v_7.2 channels, we compared the dynamics of the open channel in PIP_2_-containing lipid bilayers with a control simulation performed in the absence of PIP_2_ lipids. The most significant PIP_2_-induced conformational change was observed in the cytoplasmic helices A and B (Fig. 8a-b). The effect was quantified by calculating the orientation of the helical pair with respect to the membrane normal (Fig. 8c). In a PIP_2_-free lipid bilayer, these helices fluctuate around their initial position (θ= 83.8 ± 13.2°) and remain in a largely solvent-exposed conformation (Fig. 8b-c). In the presence of PIP_2_, the helices adopt a conformation where they interact directly with the lipid bilayer (θ= 95.5 ± 3.7°)(Fig. 8a, c). This large-scale conformational change also forms a pathway for PIP_2_ to move along Helix-B and ultimately bind to Site-O4 (Fig. 8d, Supplementary Movie 4). Since R353 in the AB linker acts as an initial anchor point for PIP_2_ binding (Supplementary Movies 1-3), we next tested if a charge-neutralizing mutation at this residue (R353Q) affects the PIP_2_-induced conformational change. We found that the R353Q mutation moves the helices back toward a more solvent-exposed conformation (θ= 86.7 ± 1.9°) (Fig. 8c).

**Figure 8.**
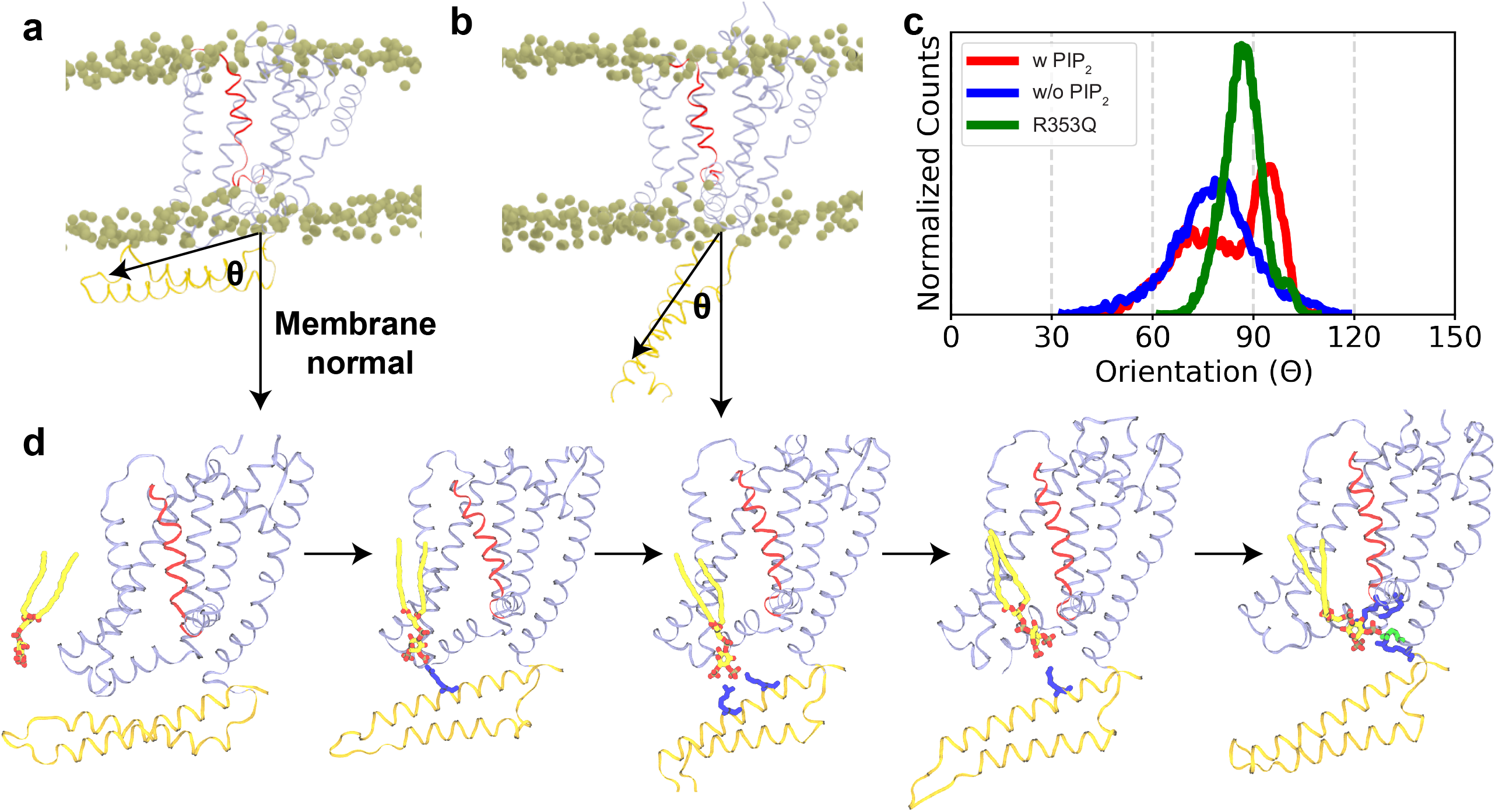
PIP_2_-mediated conformational changes of helices A and B in open K_v_7.2 channel. **(a)** Snapshot of a representative K_v_7.2 monomer in PIP_2_-containing membranes. **(b)** Snapshot of a representative K_v_7.2 monomer in PIP_2_-free membranes. **(c)** Orientation angle of helices A and B with respect to the membrane normal. The distribution of the orientation angle of helices A and B in WT K_v_7.2 channel over the last 200 ns of all the simulations in membranes with (w/) or without (w/o) PIP_2_ as well as that of mutant K_v_7.2-R353Q channel in membranes with PIP_2_. In PIP_2_-free membranes, the helical pair of the WT channel adopts a largely solvent-exposed conformation. Addition of PIP_2_ induces a drastic conformational change in these helices from a solvent-exposed conformation to a bilayer-interacting conformation. However, the R353Q mutation in the AB linker moves the helices closer to a solvent-exposed conformation even in the presence of PIP_2_. (**d**) Large-scale conformational change in the cytoplasmic helices provides a pathway for PIP_2_ movement along Helix-B to its ultimate localization at Site-O4. All the phosphorus atoms of the bilayer are shown in vdW (van der Waals) representation. K_v_7.2 protein is shown in ribbon representation with the S4 in red, helices A and B in brown, and the rest of the protein in ice blue.

## Discussion

### Identification of novel PIP_2_ binding sites in open and closed K_v_7.2 channels

Neuronal K_v_7 channels are known as the “M-channels” due to their inhibition by the activation of M1 and M3 muscarinic acetylcholine receptors ^13^. PIP_2_ hydrolysis underlies current inhibition of M-channels, bringing attention to K_v_7-PIP_2_ interaction ^6,20^. Increasing the PIP_2_ level enhances current density and the open probability of K_v_7.2 channels and induces a hyperpolarized shift in their voltage-dependence _6,14,25,28,48_. However, the detailed mechanism underlying PIP_2_-dependent regulation of neuronal K_v_7 channel remains unclear. In this study, we address this knowledge gap by identifying PIP_2_ interaction sites in both open and closed K_v_7.2 channels.

Our all-atom MD simulations have revealed PIP_2_ localization to 3 sites at the periphery of the VSD in the closed K_v_7.2 channel, whereas PIP_2_ binds to 4 distinct sites at the VSD, the pore domain, and intracellular helices in the open channel (Fig. 1, Supplementary Figs. S1-S2). The common PIP_2_ binding domains in both open and closed channels are the intracellular N-terminal tail, the S2-S3 and AB linkers. Importantly, PIP_2_ binding in the open channel is coordinated by multiple basic residues from different functional domains of K_v_7.2.

Our identification of the S2-S3 and S4-S5 linkers as PIP_2_ binding domains is consistent with a previous simulation study reported by Zhang et al ^50^. However, our study has identified many more PIP_2_ binding regions in K_v_7.2 channel compared to Zhang et al. These include the intracellular N-terminal tail, the distal ends of the S4 and S6 as well as pre-Helix-A, Helix-A, the AB linker, Helix-B, and the BC linker in the intracellular C-terminal tail. The difference could be attributed to structural templates used in each study. Our homology model of K_v_7.2 was based on the cryo-EM structure of K_v_7.1 containing cytoplasmic helices A-C and a part of intracellular N-terminal tail ^10^. In contrast, Zhang et al modeled the K_v_7.2 transmembrane domains only (residues 95–337) based on the crystal structures of K_v_1.2 and an activated bacterial K^+^ channel KcsA ^50^. In the following sections, we discuss the functional implications of the additional PIP_2_ binding sites, which we have identified in this study.

### PIP_2_ binding couples the voltage-sensor and the pore domain of K_v_7.2 channels

Our MD simulations have identified a novel PIP_2_-interacting interface in the open K_v_7.2 channel, Site-O4, which is comprised of the distal ends of S4 and S6, the S4-S5 linker, and pre-Helix-A. We show that R325 in pre-Helix-A interacts with PIP_2_ (Fig. 2), and its charge-neutralizing mutation, R325Q, abolishes this interaction (Fig. 3), basal current, and the sensitivity to PIP_2_ (Fig. 4,6). Severe reduction in current expression and PIP_2_ sensitivity has also been reported in K_v_7.2 channels containing recurrent EE variant R325G ^28^ which causes drug-resistant seizures, neurodevelopmental delay, and intellectual disability ^18,51^. The R325 is the first residue in a “RQKH” motif conserved in all K_v_7 subunits ^3,22,28^. This motif in K_v_7.1 undergoes a PIP_2_-induced conformational change from an unstructured loop to a helix ^22^, further supporting that PIP_2_ binding to R325 is involved in activation of K_v_7.2 channels.

In Site-O4, PIP_2_ also interacts with R214 in distal S4 and K219 in the S4-S5 linker in the open channel (Fig. 1, 2). Although the S4-S5 linker was previously reported to mediate PIP_2_ modulation of K_v_7.1-K_v_7.3 ^3,21,50,52^, the identification of R214 as a PIP_2_ binding site in K_v_7.2 is a novel finding. R214 is also the target of two epilepsy mutations ^53-55^, supporting its functional importance. The charge-neutralizing mutations R214Q and K219N reduce PIP_2_ interactions with the mutated residues (Fig. 3) and channel sensitivity to PIP_2_ depletion (Fig. 6). K219N, but not R214Q, induces a depolarizing shift in voltage dependence (Fig. 4) similar to R243A and K248A in the corresponding residue in K_v_7.3 ^27,52^. However, the double R214Q/K219N mutation further shifts the activation curve to a depolarized potential and induces less sensitivity to PIP_2_ depletion compared to single mutants (Figs. 4, 6), demonstrating the synergetic effects. These findings indicate that PIP_2_ interaction with both the distal S4 and the S4-S5 linker is needed for robust voltage-dependent activation of K_v_7.2 channels.

Previous studies proposed the role of PIP_2_ in coupling of the VSD activation to pore opening in K_v_7.1 and K_v_7.3 ^10,21,22,52,56^. In K_v_7.2 channels, we observe that PIP_2_ binding to Site-O4 forms an allosteric network of interactions, leading to a correlated motion of the main voltage sensor S4, the S4-S5 linker, the gate S6, and pre-Helix-A (Fig. 7), suggestive of the coupling of the VSD to the pore domain. Charge neutralizing mutations (R214Q, K219N, or R325Q) induce PIP_2_ dissociation from the mutated residues (Fig. 3) and disrupt the coordinated movements of the VSD and the pore domain (Fig. 7), indicative of their decoupling. To the best of our knowledge, this is the first study to provide an atomic-level structural basis for the allosteric role of PIP_2_ in voltage-dependent activation of neuronal K_v_7.2 channels.

### PIP_2_ binding induces conformational change important for opening K_v_7.2 channels

Our MD simulations have identified Site-O2 comprised of the K552-R553-K554 motif in Helix-B and with R560 in the BC linker as a PIP_2_ binding site in the open state (Fig. 2). We have previously shown that current potentiation induced by increasing PIP_2_ level is abolished by EE mutations K552T, R553L, and R560W ^14,25^ but enhanced by K554N ^25^, highlighting the role of these basic residues in PIP_2_ modulation of K_v_7.2 channels ^14,25^. The corresponding basic residues in Helix-B of K_v_7.1 bind to PIP_2_ *in vitro* ^23^ but those in K_v_7.3 do not affect its sensitivity to PIP_2_ depletion ^27^, suggesting the subunit-specific difference of Helix-B in mediating PIP_2_ binding.

Our simulations also demonstrate that PIP_2_ interacts with Site-O1 formed by F346, Y347, and R353 at the beginning of the AB linker and with K162 and R165 in the S2-S3 linker (Fig. 2). Due to the presence of a lysine residue (K162) and two aromatic residues (F346 and Y347), this site fulfills the requirement for a canonical PIP_2_ binding site ^57^. Consistent with the conserved sequence of the S2-S3 linker among K_v_7 subunits ^3^, PIP_2_ binding to this linker has also been reported in K_v_7.1 ^3,21,22,58^ and K_v_7.3 ^27,52^, suggesting the role of this linker in PIP_2_ modulation of K_v_7 channels.

Although a “cationic cluster” (K452/R459/K461) in the distal region of the AB linker is implicated in PIP_2_ modulation of K_v_7.2 current ^26^, the charge-neutralizing mutation of R353 in the proximal region of this linker reduces basal current expression of K_v_7.2 channels but does not alter their sensitivity to PIP_2_ (Fig. 4-6). However, combination of R353Q with R214Q and K219N in Site-O4 significantly reduces the channel’s sensitivity to PIP_2_ (Fig. 4, 6), suggesting that R353 modulates K_v_7.2 channels by coordinating PIP_2_ binding with other residues. Indeed, R353 serves as an initial anchor point for PIP_2_ binding to Site-O1 and Site-O4 (Supplementary Movies 1-4). Furthermore, PIP_2_ induces the conformational change of helices A and B from the largely solvent-exposed conformation to a bilayer-interacting conformation, whereas the R353Q mutation attenuates this effect (Fig. 8), suggesting that PIP_2_ binding to R353 contributes to this conformational change.

The conformational change in K_v_7.2 is different from that observed in the PIP_2_-bound K_v_7.1 ^22,59^ which lacks the analogous arginine residue and shows low sequence homology to the AB linker of K_v_7.2. K_v_7.3, on the other hand, has the analogous arginine residue in the AB linker with fairly well-conserved sequence, suggesting that K_v_7.3 may adopt a similar conformation as K_v_7.2 in the presence of PIP_2_. We propose that PIP_2_ binding to R353 affects the opening of K_v_7.2 channels by inducing a bilayer-interacting conformation of the helical pair and facilitating the movement of PIP_2_ along Helix-B and its binding to distal Helix-B (Site-O2). Subsequent interaction with other basic residues in Site-O1 and Site-O4 will induce the transition from the closed to the open state by coordinating allosteric movement of the VSD and the pore domain. R353 is also the target of three epilepsy mutations (ClinVar Database, NCBI) ^60^, further supporting the role of R353 in PIP_2_ modulation of K_v_7.2 channels.

### Physiological impact of K_v_7.2-PIP_2_ interaction on neuronal excitability

Neuronal K_v_7 channels are highly enriched in the axonal surface where they regulate the AP firing threshold, frequency and shape ^13,61^, whereas dendritic K_v_7 currents increase the threshold of Ca^2+^ spike initiation ^62^. Heterozygous deletion of *KCNQ2* gene in mice leads to hippocampal hyperexcitability and increased seizure propensity ^63,64^. Consistent with their critical role in inhibiting neuronal excitability, > 400 BFNE and EE mutations are found in *KCNQ2* ^17^. Computational algorithms have identified the S4, the pore loop, the S6, pre-Helix-A, Helix B, and the BC linker as hotspots for EE mutations ^14,65^, whereas BFNE mutations are enriched in the S2-S3 linker ^65^. The overlap between epilepsy mutation hotspots and the PIP_2_ binding domains identified in our study underscores the critical role of PIP_2_ in the pathophysiological mechanism underlying *KCNQ2*-related epilepsy.

Current anti-epileptic drugs are ineffective in treating many epilepsy patients with K_v_7.2 EE variants _18,19,66_. M-current inhibition upon PIP_2_ depletion results in neuronal hyperexcitability ^67^, and impaired PIP_2_ sensitivity of K_v_7.2 channels is associated with EE variants ^14,25,28^. Retigabine (INN; USAN ezogabine) is a selective agonist of K_v_7.2-K_v_7.5 channels, but not K_v_7.1 channel ^68,69^. Retigabine suppresses seizures in animal models and humans ^68,69^, although it has been discontinued as an anti-epileptic drug due to adverse side effects ^70^. However, it may be effective in opening EE mutant channels with impaired PIP_2_ binding because it stabilizes the open state of K_V_7.2 and K_V_7.3 channels by binding to a hydrophobic pocket near the gate ^71^. Alternatively, strengthening PIP_2_-K_v_7.2 interaction may increase K_v_7 current. For example, zinc pyrithione can rescue M-current in hippocampal neurons following PIP_2_ depletion by competing with PIP_2_ for K_v_7.2 activation ^72^. Recently, a compound, CP1, is shown to substitute PIP_2_ for the VSD-pore coupling in K_v_7.1 channel, and to a less extent, K_v_7.2 and K_v_7.2/K_v_7.3 channels ^73^, suggesting its potential to inhibit neuronal hyperexcitability. Our in-depth investigation of PIP_2_-K_v_7.2 interaction may provide the foundation to explore a new class of therapeutics for epilepsy that can control PIP_2_ modulation of neuronal K_v_7 channels.

## Methods

### Structural models of open and closed states of K_v_7.2

The open and closed conformations of K_v_7.2 channel used in the molecular dynamics (MD) simulations were modeled following the procedure described in a recent study ^14^. Briefly, the closed conformation of the channel was modeled based on the cryo-EM structure of K_v_7.1 (PDB ID: 5VMS) ^10^. Multiple sequence alignment of the target template and K_v_7.2 was performed using TCoffee web server (https://www.ebi.ac.uk/Tools/msa/tcoffee/). After the alignment, the homology model of the closed state was built with MODELLER ^74^. Our final model contains residues from 74-363 and 537-594. The open conformation of K_v_7.2 was then constructed by performing non-equilibrium, driven MD simulations. We performed a 20-ns targeted MD (TMD) ^75^ simulation during which the closed structure was driven towards an open form, while embedded in a lipid bilayer containing 1-palmitoyl-2-oleoyl-*sn*-glycero-3-phosphatidylcholine (POPC) and 2.2% 1-palmitoyl-2-oleoyl-*sn*-glycero-3-phosphatidylinositol 4,5-bisphosphate (PIP_2_) lipids. The target of the TMD simulation was selected to be the highly homologous K_v_1.2/K_v_2.1 channel in an open-state conformation (PDB ID: 2R9R) ^76^. As major structural changes occur in the pore region of the channel, we applied a restraint (*k* = 250 kcal/mol/Å^2^; only C*α* atoms were driven) on the S4-S5 and S6 helices of each monomer to drive it towards the target open state. The structural stability of the open and closed conformations of K_v_7.2 was evaluated by monitoring the degree of opening of the states, calculated by the number of water molecules in the pore helix and the selectivity filter during 400-ns MD runs in explicit lipid bilayers ^14^. During these simulations we consistently observed stable and a higher number of water molecules in the open state of the channel, as compared to the closed state^14^.

### PIP_2_-binding simulations

The open and closed conformations of K_v_7.2 were embedded in multiple independently generated POPC membranes with or without 2.2% PIP_2_ lipids (Table 1). All the membranes were constructed using the Membrane Builder module in CHARMM-GUI ^77^, and the initial placement of PIP_2_ lipids was intentionally varied in each membrane. All PIP_2_ lipid molecules were initially at least 15 Å away from the protein. The systems were solvated using TIP3P water and buffered at 150 mM KCl to neutralize. The final simulation systems consisted of ∼300,000 atoms. All the simulations were performed in the absence of calmodulin. To further examine the specific lipid-protein interactions captured in the modeled closed K_v_7.2, we also performed additional lipid-binding simulations of a recent cryo-EM structure of K_v_7.2 (PDB ID: 7CR3) after removing calmodulin ^44^. All the missing loops were constructed using MODELLER ^74^ and the entire tetrameric structure was embedded in an explicit lipid bilayer containing POPC and 2.2% of PIP_2_ lipids. Three independent 500-ns simulations were performed on this system.

### Molecular dynamics simulation protocols

All the simulations were performed under periodic boundary conditions using NAMD2 ^78,79^ and CHARMM36m force field parameters ^80,81^ for protein and lipid. During the initial equilibration, the protein’s backbone atoms were harmonically restrained to their initial positions with a force constant of *k* = 1 kcal/mol/Å^2^. The restraints were released at the start of the production run. All the non-bonded forces were calculated with a cutoff of 12 Å and a switching distance of 10 Å. Long-range electrostatic forces were calculated using the particle mesh Ewald (PME) method ^82^. A Langevin thermostat using γ = 1 ps^-1^ was used to maintain the system temperature at 310 K. The pressure was maintained at 1 bar using a Nosé Hoover Langevin piston method ^83^. During pressure control, the simulation box was allowed to fluctuate in all the dimensions with constant ratio in the *xy* (lipid bilayer) plane. An integration time step of 2 fs was used in all the simulations.

### Simulation Analysis

To characterize lipid-protein interactions and potential lipid binding sites, occupancy maps of the PIP_2_ headgroup were calculated using the VOLMAP plugin in VMD ^84^. Based on our previous experience ^35^, a 4-Å heavy-atom distance cutoff was chosen to define contacts between the phosphate groups of PIP_2_ lipids and protein residues, while a 3.5-Å cutoff was used to define contacts with the hydroxyl groups of PIP_2_. The role of PIP_2_ lipids in stabilizing the open conformation of K_v_7.2 was determined by performing dynamical network analysis using the NETWORK-VIEW plugin ^85^ in VMD. In a network, all Cα atoms are defined as nodes connected by edges if they are within 4.5 Å of each other for at least 75% of the MD trajectory. Pearson correlation was used to define the communities (the set of residues that move in concert) in the network.

### DNA construct and mutagenesis

Plasmid pcDNA3 carrying *KCNQ2* cDNA (GenBank: Y15065.1) encoding K_v_7.2 (GenBank: CAA 75348.1) was previously described ^14,25,86,87^. This short isoform of K_v_7.2 lacks 2 exons compared to the reference K_v_7.2 sequence (GenBank: NP_742105.1). Our lab has previously shown that currents through this K_v_7.2 variant can be potentiated by PIP_2_ increase and are sensitive to Dr-VSP activation ^14,25^. Plasmid pIRES-dsRed-PIPKI*γ*90 ^88^ was a kind gift from Dr. Anastasios Tzingounis (University of Connecticut). Selected mutations (R214Q, K219N, R325Q, R353Q, R214Q/K219N double mutation, R214Q/K219N/R353Q triple mutation) were generate using the Quik Change II XL Site-Directed Mutagenesis Kit (Agilent) and verified by sequencing the entire cDNA construct. The following primers were used for mutagenesis: R214Q (sense- ^5’^GAT CCG CAT GGA CCG G**<u>CA G</u>**GG AGG CAC CTG G^3’^, antisense- ^5’^CCA GGT GCC TCC **<u>CTG</u>** CCG GTC CAT GCG GAT C^3’^), K219N (sense- ^5’^ GGA GGC ACC TGG **<u>AAC</u>** CTG CTG GGC TCT GTG^3^’, antisense- ^5’^CAC AGA GCC CAG CAG **<u>GTT</u>** CCA GGT GCC TCC^3’^), R325Q (sense- ^5’^GGT TCA GGA GCA GCA C**<u>CA G</u>**CA GAA GCA CTT TGA GAA G^3’^, antisense- ^5’^CTT CTC AAA GTG CTT CTG **<u>CTG</u>** GTG CTG CTC CTG AAC C^3’^), R353Q (sense- ^5’^GCC ACC AAC CTC TCG **<u>CAG</u>** ACA GAC CTG CAC TCC^3’^, antisense- ^5’^GGA GTG CAG GTC TGT **<u>CTG</u>** CGA GAG GTT GGT GGC^3’^).

### Electrophysiology

Whole cell patch clamp recording was performed in Chinese ovary cells (CHO hm1) at room temperature (22-24°C) as described ^14,25^. Cells were plated on 12 mm glass coverslips (Warner Instrument, 1 ⨯ 10^5^ cells per coverslip) treated with poly D-lysine (0.1 mg/mL) (Sigma-Aldrich). To express K_v_7.2 channels and PIP5K, cells were transfected with pEGFPN1 (0.2 μg), pIRES-dsRed-PIPKI*γ*90 (0.45 μg) and pcDNA3- K_v_7.2 WT or mutant (0.45 μg). For the control experiment of PIP5K, cells were transfected with pEGFPN1 (0.65 μg) and pcDNA3-K_v_7.2 WT or mutant (0.45 μg). At 24-48 hrs post-transfection, GFP-positive cells were recorded in extracellular solution containing (mM): 138 NaCl, 5.4 KCl, 2 CaCl_2_, 1 MgCl_2_, 10 D-glucose, and 10 HEPES (pH 7.4, 307-312 mOsm). Patch pipettes (3-4 MΩ) were filled with intracellular solution containing (mM): 140 KCl, 2 MgCl_2_, 10 EGTA, 10 HEPES, 5 Mg-ATP (pH 7.4 with KOH, 290-300 mOsm).

To record K^+^ currents, cells were held at -80 mV. Currents were evoked by depolarization for 1.5 s from -100 mV to +20 mV in 10-mV increments, followed by a step to 0 mV for 300 ms. Leak-subtracted current densities (pA/pF), normalized conductance and channel biophysical properties were calculated as described ^25^. Briefly, leak current was defined as non-voltage-dependent current through GFP-transfected CHO hm1 cells. Current density was calculated by dividing leak-subtracted current (pA) by capacitance (pF). V_1/2_ and the slope factor (*k*) were calculated by fitting the points of G/G_max_ to a Boltzmann equation in the following form: G/G_max_=1/{1+exp(V_1/2_-V)/*k*}.

To examine the decline of K_v_7.2 current upon activation of Dr-VSP, CHO hm1 cells were transfected with pDrVSP-IRES2-EGFP (0.5 μg) and pcDNA3-K_v_7.2 WT or mutant (0.5 μg). The pDrVSP-IRES2-EGFP plasmid was a gift from Yasushi Okamura (Addgene plasmid # 80333). Voltage-clamp recording of K_v_7.2 current upon depolarization-induced Dr-VSP activation was performed as described ^89^ with an external solution containing 144 mM NaCl, 5 mM KCl, 2 mM CaCl_2_, 0.5 mM MgCl_2_, 10 mM glucose and 10 mM HEPES (pH 7.4). Patch pipettes (3 – 4 MΩ) were filled with intracellular solution containing 135 mM potassium aspartate, 2 mM MgCl_2_, 1 mM EGTA, 0.1 mM CaCl_2_, 4 mM ATP, 0.1 mM GTP and 10 mM HEPES (pH 7.2). Cells were held at -70 mV and 10 s step depolarizations were applied in 20-mV steps from -20 to +100 mV with 2 min inter-step intervals to allow PIP_2_ regeneration. The extent of K_v_7.2 current decay upon Dr-VSP activation during 10 s depolarization was measured as the ratio of current at 10 s over peak current at each voltage step.

### Western blot

CHO hm1 cells were plated on 35 mm tissue culture dishes (Corning, 3 × 10^5^ cells per well). Next day, cells were transfected with pcDNA3-K_v_7.2 WT or mutant (0.5 μg) using FuGENE6 transfection reagent (Promega). At 24 hrs post-transfection, cells were lysed, and lysates were analyzed by immunoblotting as previously described ^14^. Briefly, cells were collected by cell scraper in 300 μL ice-cold lysis buffer containing (mM): 150 NaCl, 50 Tris, 2 EGTA, 1 EDTA, 1% Triton-X, 0.5% deoxycholic acid (pH 7.4), supplemented with Halt protease inhibitor cocktail (Thermo Fisher Scientific). Lysates were harvested by 15 min incubation on ice, followed by 15 min centrifugation at 14,000 x g in 4°C. Samples were mixed with SDS sample buffer in 1:5 ratio and heated at 75°C for 30 min. The SDS sample buffer contained (mM): 75 Tris, 50 TCEP, 0.5 EDTA, 10% SDS, 12.5% glycerol, 0.5 mg/mL Bromophenol Blue. The samples were then run on 4%-20% gradient SDS-PAGE gels (Bio-Rad) and transferred to a methanol-treated polyvinyl difluoride (PVDF) membrane (Millipore). The membranes were blocked with blocking buffer (5% nonfat milk/0.1% Tween-20 in Tris-buffered saline/TBS containing 150 mM NaCl, 50 mM Tris, pH 7.5) for 1 hr followed by overnight incubation of primary antibodies in washing buffer (1% nonfat milk/0.1% Tween-20 in TBS) in 4°C. After 1 hr incubation with horse radish peroxidase (HRP)-conjugated secondary antibodies in washing buffer, membranes were treated with Enhanced Chemiluminescence substrate (ECL, Thermo Fisher Scientific, #32106) and immediately imaged with the iBright CL1000 imaging system (Thermo Fisher Scientific). Acquired images were analyzed using ImageJ software (National Institutes of Health). GAPDH, *α*-tubulin and β-tubulin were used as loading controls. Background-subtracted intensities of each immunoblot band was measured and the K_v_7.2/loading control ratio of WT or mutants was computed. The K_v_7.2/loading control of WT was used as 100% and mutants were normalized to WT as described ^14^. Antibodies used include anti-GAPDH (Cell Signaling #2118, 1:1000 dilution), anti-*α*-tubulin (Cell Signaling #2144, 1:1000 dilution), anti-β-tubulin (Cell Signaling #2146, 1:1000), anti-K_v_7.2 (Neuromab, N26A/23, 1:200 dilution), donkey anti-rabbit and donkey anti-mouse HRP secondary antibodies (The Jackson Laboratory, 711-035-152; 715-035-150).

### Surface biotinylation

CHO hm1 cells were plated on 60 mm culture dishes (Corning, 8 ⨯10^5^ cells per well). Next day, cells were transfected with pcDNA3-K_v_7.2 WT or mutant (0.8 μg) using FuGENE6 transfection reagent (Promega). At 24 hr post-transfection, the cells were subjected to surface biotinylation as previously described^90^. The culture dishes containing transfected cells were placed on ice and washed with 1X PBS twice. To biotinylate surface proteins, the cells were then incubated with Sulfo-NHS-SS-Biotin (1mg/mL, Pierce) in ice-cold PBS (3 mL) for 20 min. The cells were then washed with 1X PBS twice and 1X TBS once. Cells were collected using cell scraper in 400 μL ice-cold lysis buffer containing: (mM): 150 NaCl, 50 Tris, 2 EGTA, 1 EDTA, 1% Triton-X, 0.5% deoxycholic acid, supplemented with Halt protease inhibitor cocktail (Thermo Fisher Scientific). Lysates were harvested by 15 min incubation on ice, followed by 15 min centrifugation at 14,000 x g at 4°C. 40 μL lysates were saved for western blotting. The remaining 360 μL of lysates were incubated with 50% NeutraAvidin agarose beads (Pierce, 100 μL of 1:1 slurry) for overnight at 4°C to isolate biotinylated surface proteins from the lysate. After washing with the lysis buffer, biotinylated surface proteins were eluted by heating in 1x SDS sample buffer containing 50 mM TCEP at 75°C for 30 min. Eluted biotinylated surface proteins and lysates were examined by immunoblotting for K_v_7.2 and *α*-tubulin or β-tubulin as described in the previous section. ImageJ software (NIH) was used to measure background-subtracted intensity of each immunoblot band, and surface or total K_v_7.2 intensity was normalized to total *α*-tubulin or β-tubulin band intensity. To confirm that cell membranes were intact and intracellular proteins were not biotinylated during surface biotinylation, immunoblot of biotinylated surface proteins was also performed with anti-*α*-tubulin or anti-β-tubulin.

### Statistics and reproducibility

All the measurements were taken from distinct samples. All electrophysiology and immunoblotting data are reported as mean ± SEM. Origin 9.1 (OriginLab, Inc) was used for Student’s t-test and one-way ANOVA with post-hoc Fisher’s multiple comparison tests. Specifically, one-way ANOVA with post-hoc Fisher’s test was used for surface biotinylation and electrophysiology figures with the exception that Student’s unpaired t-test was used when comparing the results before and after PIP5K cotransfection in Supplementary Fig. 4a-b. Listed sample sizes of electrophysiology indicate number of cells successfully recorded, while the sample sizes of surface biotinylation studies represent the numbers of experimental replica. Statistical significance was assessed at a priori value (p) < 0.05.

## Data Availability

The data that support the findings of this study are available from the corresponding authors upon reasonable request. All relevant accession codes are provided.

## Code Availability

Simulation trajectories were collected using the simulation program NAMD. Visualization and analysis were performed using VMD and Python. All of these software packages are publicly available.

## Acknowledgments

This research was supported by the National Institutes of Health under awards R01-GM123455 and P41-GM104601 from the National Institute of General Medical Sciences (to E.T.) and R01-NS083402 and R01-NS097610 from the National Institute of Neurological Disorders and Stroke (to H.J.C.). S.P. acknowledges receiving support from the Beckman Institute Graduate Fellowship. We would like to also acknowledge the computing resources provided by Blue Waters of National Center for Supercomputing Applications (to E.T.) and by Extreme Science and Engineering Discovery Environment (XSEDE award MCA06N060 to E.T.) and TACC Frontera. We would like to thank Tao Jiang for her assistance with simulations on Frontera.

## REFERENCE

1 Di Paolo, G. & De Camilli, P. Phosphoinositides in cell regulation and membrane dynamics. Nature 443, 651–657, doi:10.1038/nature05185 (2006).

2 Suh, B. C. & Hille, B. PIP2 is a necessary cofactor for ion channel function: how and why? Annu Rev Biophys 37, 175–195, doi:10.1146/annurev.biophys.37.032807.125859 (2008).

3 Zaydman, M. A. & Cui, J. PIP2 regulation of KCNQ channels: biophysical and molecular mechanisms for lipid modulation of voltage-dependent gating. Front Physiol 5, 195, doi:10.3389/fphys.2014.00195 (2014).

4 Huang, C. L., Feng, S. & Hilgemann, D. W. Direct activation of inward rectifier potassium channels by PIP2 and its stabilization by Gbetagamma. Nature 391, 803–806, doi:10.1038/35882 (1998).

5 Rodriguez-Menchaca, A. A., Adney, S. K., Zhou, L. & Logothetis, D. E. Dual Regulation of Voltage-Sensitive Ion Channels by PIP(2). Front Pharmacol 3, 170, doi:10.3389/fphar.2012.00170 (2012).

6 Zhang, H. et al.. PIP(2) activates KCNQ channels, and its hydrolysis underlies receptor-mediated inhibition of M currents. Neuron 37, 963–975, doi:10.1016/s0896-6273(03)00125-9 (2003).

7 Maljevic, S., Wuttke, T. V., Seebohm, G. & Lerche, H. KV7 channelopathies. Pflugers Arch 460, 277–288, doi:10.1007/s00424-010-0831-3 (2010).

8 Robbins, J. KCNQ potassium channels: physiology, pathophysiology, and pharmacology. Pharmacol Ther 90, 1–19, doi:10.1016/s0163-7258(01)00116-4 (2001).

9 Cui, J. Voltage-Dependent Gating: Novel Insights from KCNQ1 Channels. Biophys J 110, 14–25, doi:10.1016/j.bpj.2015.11.023 (2016).

10 Sun, J. & MacKinnon, R. Cryo-EM Structure of a KCNQ1/CaM Complex Reveals Insights into Congenital Long QT Syndrome. Cell 169, 1042–1050 e1049, doi:10.1016/j.cell.2017.05.019 (2017).

11 Haitin, Y. & Attali, B. The C-terminus of Kv7 channels: a multifunctional module. J Physiol 586, 1803–1810, doi:10.1113/jphysiol.2007.149187 (2008).

12 Greene, D. L. & Hoshi, N. Modulation of Kv7 channels and excitability in the brain. Cell Mol Life Sci 74, 495–508, doi:10.1007/s00018-016-2359-y (2017).

13 Brown, D. A. & Passmore, G. M. Neural KCNQ (Kv7) channels. Br J Pharmacol 156, 1185–1195, doi:10.1111/j.1476-5381.2009.00111.x (2009).

14 Zhang, J. et al. Identifying mutation hotspots reveals pathogenetic mechanisms of KCNQ2 epileptic encephalopathy. Sci Rep 10, 4756, doi:10.1038/s41598-020-61697-6 (2020).

15 Lehman, A. et al. Loss-of-Function and Gain-of-Function Mutations in KCNQ5 Cause Intellectual Disability or Epileptic Encephalopathy. Am J Hum Genet 101, 65–74, doi:10.1016/j.ajhg.2017.05.016 (2017).

16 Charlier, C. et al. A pore mutation in a novel KQT-like potassium channel gene in an idiopathic epilepsy family. Nat Genet 18, 53–55, doi:10.1038/ng0198-53 (1998).

17 Miceli, F. et al.. in GeneReviews((R)) (eds M. P. Adam et al.) (1993).

18 Weckhuysen, S. et al. KCNQ2 encephalopathy: emerging phenotype of a neonatal epileptic encephalopathy. Ann Neurol 71, 15–25, doi:10.1002/ana.22644 (2012).

19 Millichap, J. J. et al. KCNQ2 encephalopathy: Features, mutational hot spots, and ezogabine treatment of 11 patients. Neurol Genet 2, e96, doi:10.1212/NXG.0000000000000096 (2016).

20 Suh, B. C. & Hille, B. Recovery from muscarinic modulation of M current channels requires phosphatidylinositol 4,5-bisphosphate synthesis. Neuron 35, 507–520, doi:10.1016/s0896-6273(02)00790-0 (2002).

21 Zaydman, M. A. et al. Kv7.1 ion channels require a lipid to couple voltage sensing to pore opening. Proc Natl Acad Sci U S A 110, 13180–13185, doi:10.1073/pnas.1305167110 (2013).

22 Sun, J. & MacKinnon, R. Structural Basis of Human KCNQ1 Modulation and Gating. Cell 180, 340–347 e349, doi:10.1016/j.cell.2019.12.003 (2020).

23 Tobelaim, W. S. et al. Competition of calcified calmodulin N lobe and PIP2 to an LQT mutation site in Kv7.1 channel. Proc Natl Acad Sci U S A 114, E869–E878, doi:10.1073/pnas.1612622114 (2017).

24 Li, Y. et al. KCNE1 enhances phosphatidylinositol 4,5-bisphosphate (PIP2) sensitivity of IKs to modulate channel activity. Proc Natl Acad Sci U S A 108, 9095–9100, doi:10.1073/pnas.1100872108 (2011).

25 Kim, E. C. et al. Reduced axonal surface expression and phosphoinositide sensitivity in Kv7 channels disrupts their function to inhibit neuronal excitability in Kcnq2 epileptic encephalopathy. Neurobiol Dis 118, 76–93, doi:10.1016/j.nbd.2018.07.004 (2018).

26 Hernandez, C. C., Zaika, O. & Shapiro, M. S. A carboxy-terminal inter-helix linker as the site of phosphatidylinositol 4,5-bisphosphate action on Kv7 (M-type) K+ channels. J Gen Physiol 132, 361–381, doi:10.1085/jgp.200810007 (2008).

27 Choveau, F. S., De la Rosa, V., Bierbower, S. M., Hernandez, C. C. & Shapiro, M. S. Phosphatidylinositol 4,5-bisphosphate (PIP2) regulates KCNQ3 K+ channels by interacting with four cytoplasmic channel domains. J Biol Chem 293, 19411–19428, doi:10.1074/jbc.RA118.005401 (2018).

28 Soldovieri, M. V. et al. Early-onset epileptic encephalopathy caused by a reduced sensitivity of Kv7.2 potassium channels to phosphatidylinositol 4,5-bisphosphate. Sci Rep 6, 38167, doi:10.1038/srep38167 (2016).

29 Ambrosino, P. et al. Kv7.3 Compound Heterozygous Variants in Early Onset Encephalopathy Reveal Additive Contribution of C-Terminal Residues to PIP2-Dependent K+ Channel Gating. Mol Neurobiol 55, 7009–7024, doi:10.1007/s12035-018-0883-5 (2018).

30 Muller, M. P. et al. Characterization of Lipid-Protein Interactions and Lipid-Mediated Modulation of Membrane Protein Function through Molecular Simulation. Chem Rev 119, 6086–6161, doi:10.1021/acs.chemrev.8b00608 (2019).

31 Duncan, A. L., Corey, R. A. & Sansom, M. S. P. Defining how multiple lipid species interact with inward rectifier potassium (Kir2) channels. Proc Natl Acad Sci U S A 117, 7803–7813, doi:10.1073/pnas.1918387117 (2020).

32 Soubias, O. et al. Membrane surface recognition by the ASAP1 PH domain and consequences for interactions with the small GTPase Arf1. Sci Adv 6, doi:10.1126/sciadv.abd1882 (2020).

33 Miranda, W. E. et al. Lipid regulation of hERG1 channel function. Nat Commun 12, 1409, doi:10.1038/s41467-021-21681-8 (2021).

34 Kasimova, M. A., Tarek, M., Shaytan, A. K., Shaitan, K. V. & Delemotte, L. Voltage-gated ion channel modulation by lipids: insights from molecular dynamics simulations. Biochim Biophys Acta 1838, 1322–1331, doi:10.1016/j.bbamem.2014.01.024 (2014).

35 Yu, K., Jiang, T., Cui, Y., Tajkhorshid, E. & Hartzell, H. C. A network of phosphatidylinositol 4,5-bisphosphate binding sites regulates gating of the Ca2+-activated Cl-channel ANO1 (TMEM16A). Proc Natl Acad Sci U S A 116, 19952–19962, doi:10.1073/pnas.1904012116 (2019).

36 Schwake, M., Pusch, M., Kharkovets, T. & Jentsch, T. J. Surface expression and single channel properties of KCNQ2/KCNQ3, M-type K+ channels involved in epilepsy. J Biol Chem 275, 13343–13348, doi:10.1074/jbc.275.18.13343 (2000).

37 Wang, H. S. et al. KCNQ2 and KCNQ3 potassium channel subunits: molecular correlates of the M-channel. Science 282, 1890–1893, doi:10.1126/science.282.5395.1890 (1998).

38 Etxeberria, A., Santana-Castro, I., Regalado, M. P., Aivar, P. & Villarroel, A. Three mechanisms underlie KCNQ2/3 heteromeric potassium M-channel potentiation. J Neurosci 24, 9146–9152, doi:10.1523/JNEUROSCI.3194-04.2004 (2004).

39 Zaika, O., Hernandez, C. C., Bal, M., Tolstykh, G. P. & Shapiro, M. S. Determinants within the turret and pore-loop domains of KCNQ3 K+ channels governing functional activity. Biophys J 95, 5121–5137, doi:10.1529/biophysj.108.137604 (2008).

40 Soh, H., Pant, R., LoTurco, J. J. & Tzingounis, A. V. Conditional deletions of epilepsy-associated KCNQ2 and KCNQ3 channels from cerebral cortex cause differential effects on neuronal excitability. J Neurosci 34, 5311–5321, doi:10.1523/JNEUROSCI.3919-13.2014 (2014).

41 Soldovieri, M. V., Miceli, F. & Taglialatela, M. Driving with no brakes: molecular pathophysiology of Kv7 potassium channels. Physiology (Bethesda) 26, 365–376, doi:10.1152/physiol.00009.2011 (2011).

42 Nappi, P. et al. Epileptic channelopathies caused by neuronal Kv7 (KCNQ) channel dysfunction. Pflugers Arch 472, 881–898, doi:10.1007/s00424-020-02404-2 (2020).

43 Gomis-Perez, C. et al. Homomeric Kv7.2 current suppression is a common feature in KCNQ2 epileptic encephalopathy. Epilepsia 60, 139–148, doi:10.1111/epi.14609 (2019).

44 Li, X. et al. Molecular basis for ligand activation of the human KCNQ2 channel. Cell Res 31, 52–61, doi:10.1038/s41422-020-00410-8 (2021).

45 Kosenko, A. et al. Coordinated signal integration at the M-type potassium channel upon muscarinic stimulation. EMBO J 31, 3147–3156, doi:10.1038/emboj.2012.156 (2012).

46 Gamper, N., Stockand, J. D. & Shapiro, M. S. The use of Chinese hamster ovary (CHO) cells in the study of ion channels. J Pharmacol Toxicol Methods 51, 177–185, doi:10.1016/j.vascn.2004.08.008 (2005).

47 van den Bout, I. & Divecha, N. PIP5K-driven PtdIns(4,5)P2 synthesis: regulation and cellular functions. J Cell Sci 122, 3837–3850, doi:10.1242/jcs.056127 (2009).

48 Li, Y., Gamper, N., Hilgemann, D. W. & Shapiro, M. S. Regulation of Kv7 (KCNQ) K+ channel open probability by phosphatidylinositol 4,5-bisphosphate. J Neurosci 25, 9825–9835, doi:10.1523/JNEUROSCI.2597-05.2005 (2005).

49 Jensen, M. O. et al. Mechanism of voltage gating in potassium channels. Science 336, 229–233, doi:10.1126/science.1216533 (2012).

50 Zhang, Q. et al. Dynamic PIP2 interactions with voltage sensor elements contribute to KCNQ2 channel gating. Proc Natl Acad Sci U S A 110, 20093–20098, doi:10.1073/pnas.1312483110 (2013).

51 Pisano, T. et al. Early and effective treatment of KCNQ2 encephalopathy. Epilepsia 56, 685–691, doi:10.1111/epi.12984 (2015).

52 Kim, R. Y., Pless, S. A. & Kurata, H. T. PIP2 mediates functional coupling and pharmacology of neuronal KCNQ channels. Proc Natl Acad Sci U S A 114, E9702–E9711, doi:10.1073/pnas.1705802114 (2017).

53 Fang, Z. X. et al. KCNQ2 related early-onset epileptic encephalopathies in Chinese children. J Neurol 266, 2224–2232, doi:10.1007/s00415-019-09404-y (2019).

54 Miraglia del Giudice, E. et al. Benign familial neonatal convulsions (BFNC) resulting from mutation of the KCNQ2 voltage sensor. Eur J Hum Genet 8, 994–997, doi:10.1038/sj.ejhg.5200570 (2000).

55 Castaldo, P. et al. Benign familial neonatal convulsions caused by altered gating of KCNQ2/KCNQ3 potassium channels. J Neurosci 22, RC199 (2002).

56 Hou, P. et al. Two-stage electro-mechanical coupling of a KV channel in voltage-dependent activation. Nat Commun 11, 676, doi:10.1038/s41467-020-14406-w (2020).

57 Slesinger, P. A. & Wickman, K. Structure to Function of G Protein-Gated Inwardly Rectifying (GIRK) Channels. 1 edn, Vol. 123 (Academic Press, 2015).

58 Eckey, K. et al. Novel Kv7.1-phosphatidylinositol 4,5-bisphosphate interaction sites uncovered by charge neutralization scanning. J Biol Chem 289, 22749–22758, doi:10.1074/jbc.M114.589796 (2014).

59 Kang, P. W. et al. Calmodulin acts as a state-dependent switch to control a cardiac potassium channel opening. Sci Adv 6, doi:10.1126/sciadv.abd6798 (2020).

60 Milh, M. et al. Variable clinical expression in patients with mosaicism for KCNQ2 mutations. Am J Med Genet A 167A, 2314–2318, doi:10.1002/ajmg.a.37152 (2015).

61 Shah, M. M., Migliore, M., Valencia, I., Cooper, E. C. & Brown, D. A. Functional significance of axonal Kv7 channels in hippocampal pyramidal neurons. Proc Natl Acad Sci U S A 105, 7869–7874, doi:10.1073/pnas.0802805105 (2008).

62 Yue, C. & Yaari, Y. Axo-somatic and apical dendritic Kv7/M channels differentially regulate the intrinsic excitability of adult rat CA1 pyramidal cells. J Neurophysiol 95, 3480–3495, doi:10.1152/jn.01333.2005 (2006).

63 Kim, E. C. et al. Heterozygous loss of epilepsy gene KCNQ2 alters social, repetitive and exploratory behaviors. Genes Brain Behav 19, e12599, doi:10.1111/gbb.12599 (2020).

64 Watanabe, H. et al. Disruption of the epilepsy KCNQ2 gene results in neural hyperexcitability. J Neurochem 75, 28–33, doi:10.1046/j.1471-4159.2000.0750028.x (2000).

65 Goto, A. et al. Characteristics of KCNQ2 variants causing either benign neonatal epilepsy or developmental and epileptic encephalopathy. Epilepsia 60, 1870–1880, doi:10.1111/epi.16314 (2019).

66 Numis, A. L. et al. KCNQ2 encephalopathy: delineation of the electroclinical phenotype and treatment response. Neurology 82, 368–370, doi:10.1212/WNL.0000000000000060 (2014).

67 Delmas, P. & Brown, D. A. Pathways modulating neural KCNQ/M (Kv7) potassium channels. Nat Rev Neurosci 6, 850–862, doi:10.1038/nrn1785 (2005).

68 Blackburn-Munro, G., Dalby-Brown, W., Mirza, N. R., Mikkelsen, J. D. & Blackburn-Munro, R. E. Retigabine: chemical synthesis to clinical application. CNS Drug Rev 11, 1–20, doi:10.1111/j.1527-3458.2005.tb00033.x (2005).

69 Gunthorpe, M. J., Large, C. H. & Sankar, R. The mechanism of action of retigabine (ezogabine), a first-in-class K+ channel opener for the treatment of epilepsy. Epilepsia 53, 412–424, doi:10.1111/j.1528-1167.2011.03365.x (2012).

70 Mathias, S. V. & Abou-Khalil, B. W. Ezogabine skin discoloration is reversible after discontinuation. Epilepsy Behav Case Rep 7, 61–63, doi:10.1016/j.ebcr.2017.01.001 (2017).

71 Kim, R. Y. et al. Atomic basis for therapeutic activation of neuronal potassium channels. Nat Commun 6, 8116, doi:10.1038/ncomms9116 (2015).

72 Zhou, P. et al. Phosphatidylinositol 4,5-bisphosphate alters pharmacological selectivity for epilepsy-causing KCNQ potassium channels. Proc Natl Acad Sci U S A 110, 8726–8731, doi:10.1073/pnas.1302167110 (2013).

73 Liu, Y. et al. A PIP2 substitute mediates voltage sensor-pore coupling in KCNQ activation. Commun Biol 3, 385, doi:10.1038/s42003-020-1104-0 (2020).

74 Webb, B. & Sali, A. Comparative Protein Structure Modeling Using MODELLER. Curr Protoc Bioinformatics 54, 5 6 1–5 6 37, doi:10.1002/cpbi.3 (2016).

75 Schlitter, J., Engels, M., Krüger, P., Jacoby, E. & Wollmer, A. Targeted Molecular Dynamics Simulation of Conformational Change-Application to the T ? R Transition in Insulin. Molecular Simulation 10, 291–308, doi:10.1080/08927029308022170 (1993).

76 Long, S. B., Tao, X., Campbell, E. B. & MacKinnon, R. Atomic structure of a voltage-dependent K+ channel in a lipid membrane-like environment. Nature 450, 376–382, doi:10.1038/nature06265 (2007).

77 Jo, S., Kim, T., Iyer, V. G. & Im, W. CHARMM-GUI: a web-based graphical user interface for CHARMM. J Comput Chem 29, 1859–1865, doi:10.1002/jcc.20945 (2008).

78 Phillips, J. C. et al. Scalable molecular dynamics with NAMD. J Comput Chem 26, 1781–1802, doi:10.1002/jcc.20289 (2005).

79 Phillips, J. C. et al. Scalable molecular dynamics on CPU and GPU architectures with NAMD. J Chem Phys 153, 044130, doi:10.1063/5.0014475 (2020).

80 Best, R. B. et al. Optimization of the additive CHARMM all-atom protein force field targeting improved sampling of the backbone phi, psi and side-chain chi(1) and chi(2) dihedral angles. J Chem Theory Comput 8, 3257–3273, doi:10.1021/ct300400x (2012).

81 Klauda, J. B. et al. Update of the CHARMM all-atom additive force field for lipids: validation on six lipid types. J Phys Chem B 114, 7830–7843, doi:10.1021/jp101759q (2010).

82 Essmann, U. et al. A smooth particle mesh Ewald method. The Journal of Chemical Physics 103, 8577–8593, doi:10.1063/1.470117 (1995).

83 Martyna, G. J., Tobias, D. J. & Klein, M. L. Constant pressure molecular dynamics algorithms. The Journal of Chemical Physics 101, 4177–4189, doi:10.1063/1.467468 (1994).

84 Humphrey, W., Dalke, A. & Schulten, K. VMD: visual molecular dynamics. J Mol Graph 14, 33-38, 27-38, doi:10.1016/0263-7855(96)00018-5 (1996).

85 Sethi, A., Eargle, J., Black, A. A. & Luthey-Schulten, Z. Dynamical networks in tRNA:protein complexes. Proc Natl Acad Sci U S A 106, 6620–6625, doi:10.1073/pnas.0810961106 (2009).

86 Cavaretta, J. P. et al. Polarized axonal surface expression of neuronal KCNQ potassium channels is regulated by calmodulin interaction with KCNQ2 subunit. PLoS One 9, e103655, doi:10.1371/journal.pone.0103655 (2014).

87 Chung, H. J., Jan, Y. N. & Jan, L. Y. Polarized axonal surface expression of neuronal KCNQ channels is mediated by multiple signals in the KCNQ2 and KCNQ3 C-terminal domains. Proc Natl Acad Sci U S A 103, 8870–8875, doi:10.1073/pnas.0603376103 (2006).

88 Kim, K. S., Duignan, K. M., Hawryluk, J. M., Soh, H. & Tzingounis, A. V. The Voltage Activation of Cortical KCNQ Channels Depends on Global PIP2 Levels. Biophys J 110, 1089–1098, doi:10.1016/j.bpj.2016.01.006 (2016).

89 Kosenko, A. & Hoshi, N. A change in configuration of the calmodulin-KCNQ channel complex underlies Ca2+-dependent modulation of KCNQ channel activity. PLoS One 8, e82290, doi:10.1371/journal.pone.0082290 (2013).

90 Baculis, B. C. et al. Prolonged seizure activity causes caspase dependent cleavage and dysfunction of G-protein activated inwardly rectifying potassium channels. Sci Rep 7, 12313, doi:10.1038/s41598-017-12508-y (2017).

